# ACTOR: a latent Dirichlet model to compare expressed isoform proportions to a reference panel

**DOI:** 10.1101/856401

**Authors:** Sean D. McCabe, Andrew B. Nobel, Michael I. Love

## Abstract

The relative proportion of RNA isoforms expressed for a given gene has been associated with disease states in cancer, retinal diseases, and neurological disorders. Examination of relative isoform proportions can help determine biological mechanisms, but such analyses often require a per-gene investigation of splicing patterns. Leveraging large public datasets produced by genomic consortia as a reference, one can compare splicing patterns in a dataset of interest with those of a reference panel in which samples are divided into distinct groups (tissue of origin, disease status, etc). We propose ACTOR, a latent Dirichlet model with Dirichlet Multinomial observations to compare expressed isoform proportions in a dataset to an independent reference panel. We use a variational Bayes procedure to estimate posterior distributions for the group membership of one or more samples. Using the Genotype-Tissue Expression (GTEx) project as a reference dataset, we evaluate ACTOR on simulated and real RNA-seq datasets to determine tissue-type classifications of genes. ACTOR is publicly available as an R package at https://github.com/mccabes292/actor.

## 1 Introduction

Genes can be characterized as multiple transcripts, or isoforms, which differ in expression by their inclusion of exons and by the start and end sites of transcription (Reyes and Huber (2018)). One or more transcript of a gene may be expressed in a given sample, and the proportion of total gene expression that is attributed to each transcript can be referred to as transcript usage. Studying transcript usage across samples and datasets can provide useful information that can complement studies of total gene expression. Aberrant differential transcript usage (DTU) has been found to be associated with cancer (Climente-González *and others* (2017); Vitting-Seerup and Sandelin (2017)), retinal disease, and neurological disorders (Scotti and Swanson (2016)). Isoform-level expression has also been shown to be valuable as a predictor of *in vitro* drug response in the context of screening for cytotoxic and targeted therapies on cancer cell lines, and can can be even more predictive than gene-level expression (Safikhani *and others*, 2017). Isoforms are known to be differentially spliced across tissues (Saha *and others* (2018)) with samples from the brain, pancreas, liver, and peripheral nervous system exhibiting the most distinct alternative splicing patterns (Yeo *and others* (2004)).

A number of methods aimed at detecting DTU have been developed: DRIMSeq (Nowicka and Robinson (2016)), DEXSeq (Anders *and others* (2012)), and SUPPA2 (Trincado *and others* (2018)) are three popular methods that identify genes that are differentially spliced across one or more conditions (i.e. treatment group). DRIMSeq uses a Dirichlet Multinomial distribution to model the raw isoform expression assuming that the total gene expression per sample is a fixed quantity. Precision parameters for the Dirichlet Multinomial distribution are estimated per gene using an empirical Bayes approach, that borrows strength across genes. DEXSeq, on the other hand, utilizes a Negative Binomial distribution to model the counts per exon. It was originally targeted for identifying differential exon usage, but has been also evaluated in the context of transcript usage (Soneson *and others*, 2016; Nowicka and Robinson, 2016; Love *and others*, 2018). SUPPA2 uses biological replicates to estimate differences in isoform proportions across conditions and between biological replicates. While statistical models have been proposed for investigating splicing patterns within a collection of samples across multiple conditions, we sought to develop models for comparing isoform splicing patterns in a set of samples to patterns seen in large, publicly available genomic databases.

The generation of large genomic databases has led to the discovery of many disease associations and pathways. A primary focus of the GTEx project is detection of genetic loci that may act to modulate gene expression levels across different tissues. This research has also led to the characterization of tissue-specific networks of splicing patterns Saha *and others* (2018). The Cancer Genome Atlas (TCGA) (Weinstein *and others* (2013)) has collected data for a wide range of cancers and genomic assays to improve ways of diagnosing, treating, and preventing cancer. Large genomic databases may also serve as reference panels to gain information about smaller independent datasets. Several multi-study methods exist for the purposes of utilizing publicly available reference panels to make conclusions with independent datasets. MultiPLIER (Taroni *and others* (2019)), DARTS (Zhang *and others* (2019)), and Snaptron (Wilks *and others* (2018)) are just a few examples of these multi-study methods. MultiPLIER utilizes a transfer learning framework to aid in the identification of rare disease associations with gene expression in small datasets by borrowing information available in a reference panel. DARTS uses reference panels to increase power in the identification of differential alternative splicing in small sample datasets through the implementation of a deep neural network. Snaptron is a tool for querying and subsetting the visualization of isoform splicing patterns from over 70,000 samples from SRAv1, SRAv2, GTEx, and TCGA. IsoformSwitchAnalyzeR (Vitting-Seerup and Sandelin (2019)) uses known protein domains to label isoform switches and to identify downstream functional changes due to differential splicing. IsoformSwitchAnalyzeR utilizes the results of a differential splicing analysis and existing annotation databases. Likewise, tappAS (de la Fuente *and others* (2019)) uses annotation databases of functional domains and motifs to provide functional annotation of differential isoform analyses for a given dataset.

At the gene level, expression can be functionally characterized by known gene sets. However, there is less annotation data at the transcript level and there are fewer statistical methods designed specifically for recognizing characteristic splicing patterns. An alternative approach would be to characterize the splicing patterns in a sample, or set of samples, by comparing the patterns to those seen in a large reference panel. While some methodological comparisons have been made in the area of clustering of samples based on isoform splicing, less comparisons or method development has been made in comparing experimental data to a reference panel. We propose ACTOR, **A** latent Dirichlet model to **C**ompare expressed isoform proportions **TO** a **R**eference panel, to relate isoform splicing patterns in an experimental dataset to an independent reference panel composed of discrete sample groupings. We provide gene level interpretations of the relatedness across experimental samples and provide an improvement to standard distance based approaches. ACTOR was validated using data from the Genotype-Tissue Expression project (GTEx) () and accurately identified genes and gene classes for which isoform splicing was similar across tissues.

## 2 Distances on Isoform Proportions

We first consider the literature on unsupervised comparisons of individual samples based on their isoform splicing patterns. Johnson and Purdom (2017) examined isoform splicing data for the purposes of clustering samples in an unsupervised setting. They proposed taking a weighted sum of a per gene distance as a metric of similarity between samples. Let *X*_*ijk*_ be the *i*^*th*^ sample’s isoform expression of the *k*^*th*^ isoform for gene *j*. Then the total gene expression is the sum of the expression over all isoforms 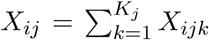. One can then calculate the proportion of the total gene expression of gene *j* which is attributable to isoform *k* as *X*_*ijk*_*/X*_*ij*_. This quantity can be calculated for all isoforms as 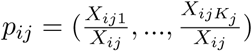. One can then define the distance between two samples (*i* and *i*^*I*^) as a weighted sum of each gene distance 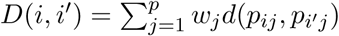. Multiple distance metrics were investigated by Johnson and Purdom including a Squared *χ*^2^, Euclidean, Jeffrey’s Divergence, Hellinger, and a log-likelihood based distance. The authors recommend a Hellinger distance for each gene given by 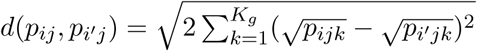 because of its superior performance in simulations and real datasets. The authors also attempted to estimate gene weights using an *l*_1_ penalty to impose sparsity, but found that it added a large amount of variability in the clustering results and did not perform well in most simulations. Gene weights were thus assigned to be constant across all genes.

A distance based approach is quite effective for the purposes of clustering samples and can also be applied to the comparison of one experimental sample with a collection of samples from a reference database. Herein, we will refer to the smaller experimental dataset of interest as the “experimental dataset” and the independent large genomic database as the “reference panel”. If the reference panel has a predefined discrete grouping of samples, distances can be used to determine which reference group a sample is most similar to. Distances can be calculated between the sample of interest and all samples in the reference panel. One could then examine the pairwise distances and identify which reference group provided a “large” number of samples with a small distance, e.g., a majority vote classifier based on the *k* closest samples in the reference panel. However, using this approach for this purpose has some limitations. A distance-based approach requires that a pairwise distance is calculated between an experimental sample and every sample in the reference dataset. In some reference panels, the number of samples can be large (∼10, 000) which creates challenges with the computational time as well as the storage capacity. Additionally, these comparisons would be performed with only one experimental sample at a time, and thus there is not a straightforward way to aggregate across experimental samples to make conclusions about the experiment as a whole. Pairwise distances do not naturally take into account the biological variability of the reference groups unless a Mahalanobis, or similar type of distance is used. A reference group could contain an outlying sample that aligns almost perfectly with an experimental sample. As a result, if conclusions are being made based on the reference sample with the smallest distance, the results may rely heavily on the presence of outliers. Distances lack a consistent scale across datasets with a varying number of genes, which may make it difficult to compare degrees of similarity of samples to reference panels across various datasets.

Another potential drawback of a distance-based approach for comparison of samples to a reference panel is that potentially interesting patterns among genes are not revealed through the analysis. For example, examining the pairwise sample distances can give insight into the general similarity of samples based on splicing patterns, however more investigation must be done to make gene level conclusions. A sample may splice similarly to only a single reference panel group for gene A, while it may splice similarly to a number of reference panel groups for gene B. It is desirable to characterize these gene-specific aspects of the splicing pattern as well as provide an aggregate view of similarity.

Due to these challenges, a probabilistic model may be more appealing, in particular one that formalizes the relationship of the sample(s) to the information provided by the reference panel. Latent Dirichlet Allocation (LDA) is a text mining model that has been applied to biological settings in the past for its ability to identify flexible mixture distributions of latent classes. We considered an extension of the LDA framework to isoform splicing, and comparison of a sample to a reference panel.

## 3 Topic Modeling - LDA

Topic modeling is a collection of statistical models aimed at discovering latent topics that occur in a set of written documents. One such model is Latent Dirichlet Allocation (LDA) (Blei *and others* (2003) and Pritchard *and others* (2000)). The use of topic modeling is becoming more prevalent in biological research with applications to the microbiome (Sankaran and Holmes (2018); Holmes *and others* (2012)) and in single cell transposase-accessible chromatin sequencing (scATAC-Seq) (González-Blas *and others* (2019)). Topic modeling for the microbiome allowed for the identification of bacterial communities with similar expression. Bacterial species may appear in multiple contexts and thus application of LDA allows for certain bacterial species to be flexibly modeled as a distribution across the latent species collections. Topic modeling for scATAC-Seq was used for the identification of cell types, enhancers, and relevant transcription factors. Another model related to the work presented here is MixDir, a Bayesian model for high dimensional clustering of samples based on high dimensional categorical data such as surveys and questionnaires (Ahlmann-Eltze and Yau (2018)). MixDir proposes a generative model similar to LDA, and employs latent classes to group similar observations, based on their responses to a moderately-sized number of multiple-choice questions (e.g. < 60 questions). Due to the count nature of isoform splicing, the generative model proposed in LDA is a good starting point for constructing a probabilistic model to characterize the similarity of a set of samples to predefined reference groups using mixtures.

Latent Dirichlet Allocation (LDA) has been proposed as a probabilistic model for text mining as well as in inferring population structure using genotype data. In the text mining application, LDA examines the frequencies of words in a large collection of documents to infer latent classes that characterize documents based on the distribution of word usage. These latent classes are often associated with a genre or topic for the documents. LDA follows a generative model that can be described per document below. First a probability vector (*θ*) for the latent classes is generated. Then for each word, a topic (*z*_*n*_) is chosen from a Multinomial distribution with the previous probability vector (*θ*). Finally, a word (*w*_*n*_) is randomly chosen from a Multinomial probability (*β*) conditioned on the topic (*z*_*n*_). This generative model is fit using a variational Bayes estimation procedure. The generative model for LDA is outlined below.

1. Choose a latent topic probability vector *θ* ∼ *Dir*(*α*)
2. For each of the *N* words *w*_*n*_
  a. Choose a topic *z*_*n*_ ∼ *Multinomial*(*θ*)
  b. Choose a word 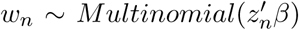, a multinomial probability specific to the latent topic

Using LDA as basis, we aim to expand on this model framework to construct a probabilistic model that compares expressed isoform splicing patterns in an experimental dataset with those in a reference panel.

## 4 ACTOR

We model the estimated isoform expression of each gene in each reference group in our reference panel using the Dirichlet Multinomial distribution. Assuming that for each gene (*g*) all samples in a particular reference group (*s*) are i.i.d. Dirichlet Multinomial, we calculate the maximum likelihood estimates for the distribution parameters, ***α***_***gs***_. These estimates can be calculated for all gene and reference group combinations, which reduces the reference panel to a more manageable size. For a sample (*j*) in the experimental dataset, we calculate a per gene (*g*) likelihood conditional on the parameter estimates coming from a specific reference panel group (*s*) as *p*(***X***_***gj***_|***α***_***gs***_). Using these likelihoods, we wish to identify latent groupings of genes that are spliced in a similar manner to our reference groups. Identifying these gene classes can be a challenging task, and topic modeling through the use of Latent Dirichlet Allocation (LDA) can provide some insight into accomplishing this task. Figure 1 provides a visualization of our method and Supplementary Table 1 provides a table of all notation for this model.

**Figure 1:**
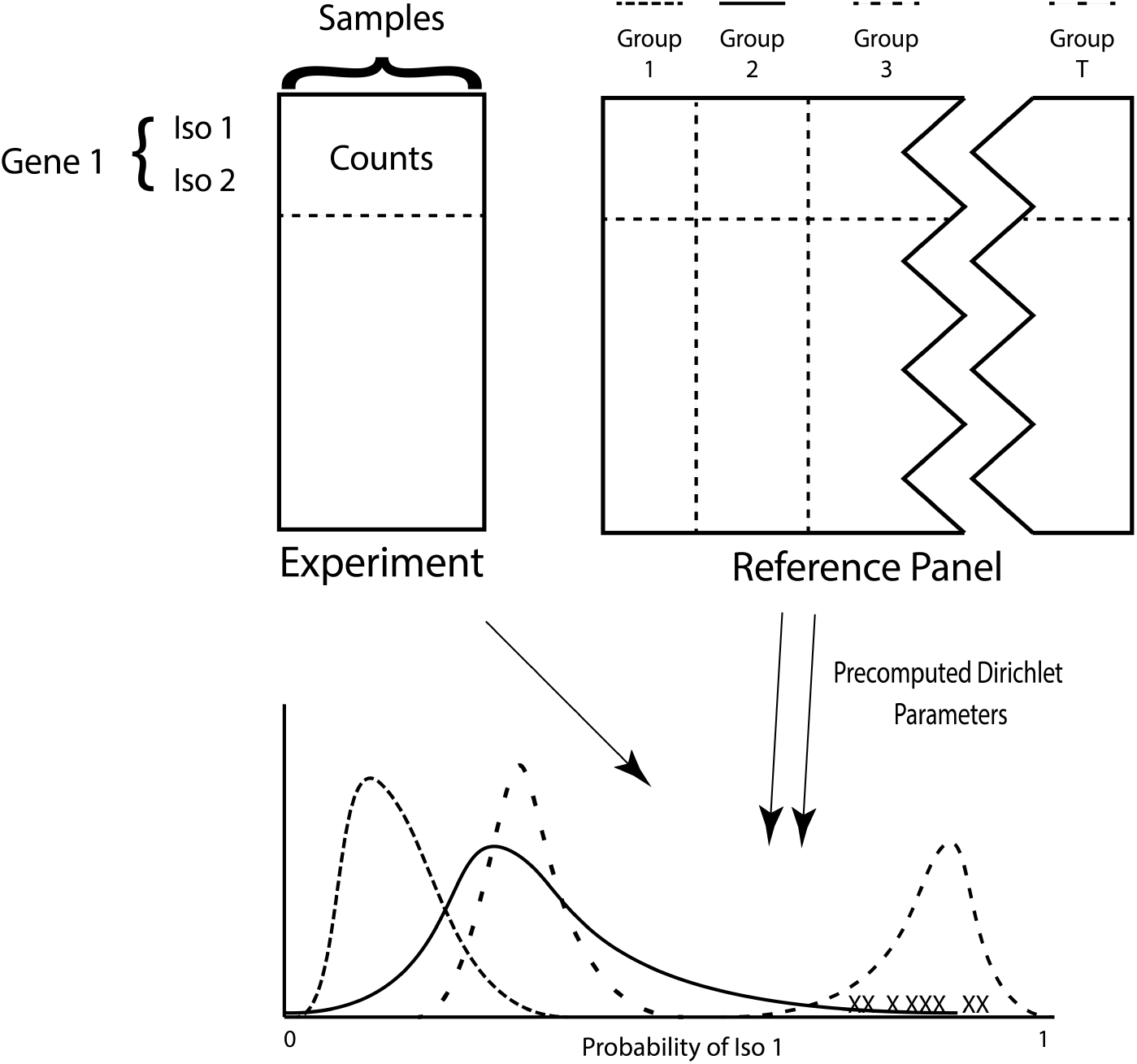
ACTOR uses precomputed Dirichlet parameters for each reference group to identify which reference group a set of experimental samples is most similar to based on isoform splicing patterns. The colored curves represent the Dirichlet distribution for each reference group. ‘X’ corresponds to an experimental sample. For visual simplicity, a one-dimensional simplex is shown here.

Expanding on the LDA framework and incorporating the pre-calculated parameter estimates for the Dirichlet Multinomial distribution, we construct a generative model identifying the mixture of reference panel groups found in the isoform splicing of an experimental dataset. Our framework is aligned with the LDA framework as follows: samples are documents, reference groups are topics, and genes are words. Additionally, we add a second latent component by modeling sets of genes that have similar reference group distributions. In this formulation, the reference group membership could be considered the latent topic from the LDA model. Dirichlet Multinomial estimates for each group in our reference panel are pre-calculated, so while the reference group assigned to a gene in our experimental dataset is a latent variable, the groups themselves are not latent, nor are the parameters describing the distribution of isoform proportion for a given group. In contrast, the topics in the text-mining LDA formulation are latent, and discovered during model fitting. This reference group assignment within the LDA model makes use of the likelihood conditional on the reference group estimates.

As mentioned earlier, it is also informative to identify classes of genes that align to the groups in the reference panel similarly. A sample could splice similarly to the reference groups across one collection of genes, while splicing similarly to a different set of reference groups in another collection of genes. The splicing could also be characterized as being unique to one reference group or as a mixture across multiple groups. We will use the term “gene class” to indicate a collection of genes that have a similar relation in the experimental samples to the reference groups. All genes within the same class have a common probability vector corresponding to the reference group from which they are generated. Supplementary figure 1 gives a toy example, with five reference panel groups and four gene classes. Three of the gene classes correspond to sets of genes that are all spliced similarly to one specific reference group. Gene class 1 aligns with reference group 1, gene class 2 aligns with reference group 2, and gene class 4 aligns with reference group 5. We also see that there could be situations in which genes are spliced similarly to two different reference groups. Gene class 3 is shown to be equally similar to reference groups 3 and 4. Therefore, we aim to estimate both reference group and gene class membership using a generative probabilistic model in a manner similar to LDA. The user prespecifies the number of gene classes for the model. As will be seen in the results, an overestimate of the number of classes will result in the model creating empty gene classes for those that are not needed.

The task addressed by our proposed model is related but nevertheless distinct from the task addressed by MixDir (Ahlmann-Eltze and Yau (2018)) in a number of aspects: while MixDir could be used to cluster thousands of genes (MixDir’s observations) based on their splicing across a moderately-sized number of samples (MixDir’s questions/features), here we are interested in grouping genes based on splicing similarity in a set of samples, compared to a reference panel. There are two relevant datasets for our task, though we can precompute distributional parameters for one of them, not just one set of relevant samples. In addition, the reference panel in our setting has known group structure (e.g. samples belonging to tissues).

ACTOR is probabilistic model is a hierarchical latent class model with the first level of latent classes being informed by the reference panel. The first step is to generate a probability vector (***γ***) of length *C* (number of gene classes) for the gene class assignment from a Dirichlet distribution with unknown parameter *ω* of length *C*. This probability vector is common for all genes. For each gene class (*l*), a probability vector of length *T* (number of reference groups) is drawn for the reference group assignment (*θ*_*l*_) from a Dirichlet distribution with unknown parameter *β. β* is of length *T* and is common across all classes. For each gene, a gene class (*c*_*g*_) is drawn from a Multinomial distribution with parameter ***γ***, which was generated in the first step. Then, using the reference group probability vector *θ*_*l*_ that corresponds with the selected gene class (*c*_*g*_), a reference group (*t*_*g*_) is drawn from the Multinomial distribution. In the formulation below, **Θ** = (***θ***_**1**_, …, ***θ***_***C***_)^′^ is a *T* × *C* matrix and ***c***_***g***_ is a vector of length *C* filled with zeros everywhere except for a 1 in the position of the gene class assignment. If ***c***_***g***_ assigns a gene to gene class *l*, then 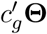 will return the probability vector for gene class *l* (*θ*_*l*_). Finally, isoform expression is generated per sample (*j*) using a Dirichlet Multinomial with parameters (***α***_***gs***_) which are the Dirichlet Multinomial parameters from the reference panel that correspond to the reference group selected by *t*_*g*_ as drawn earlier. Similarly to *c*_*g*_, *t*_*g*_ is a vector of length *T* with a 1 in the position of the selected reference group and 0 everywhere else. Likewise, we let A_*g*_ = (*α*_*g*1_, …, *α*_*gT*_)^′^ be a *T* x *I*_*g*_ matrix where *I*_*g*_ is the number of isoforms for gene *g*. Then 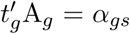 if *t*_*g*_ assigns a gene to reference group *s*. Here the Dirichlet Multinomial distribution is written as a multinomial distribution with a Dirichlet prior on the probability vector *p*_*gj*_. *p*_*gj*_ is a vector of length equal to the number of isoforms for gene *g* (*I*_*g*_). It will be seen later that ***p***_***gj***_ will not be necessary to the model and will be integrated out. Also observe that in this model the overall gene expression (*N*_*gj*_) is considered to be known and fixed. Since there is only interest in observing alignment based on isoform splicing, the generating process for gene expression is not of concern and focus is solely on how the isoform expression is allocated. The generative model is described below and a graphical representation of the model is in Figure 2a.

**Figure 2:**
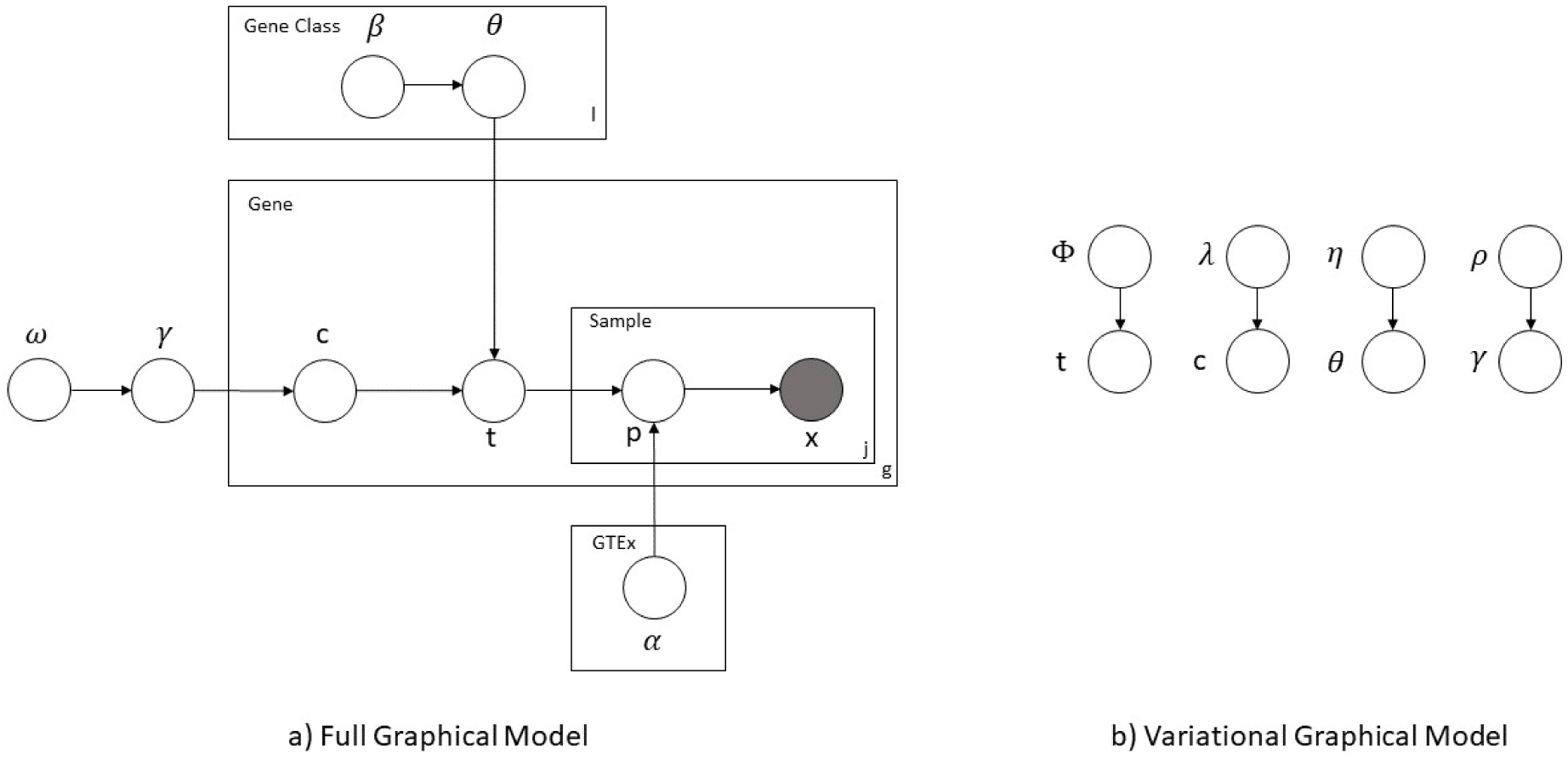
Graphical model representation of our proposed framework. Figure **a)** corresponds to the full model while Figure **b)** corresponds to the variational distribution.

> Select gene class probability vector ***γ*** ∼ *Dir*(*ω*)
>
> For each Gene Class (l)
>
> Select reference group probability vector ***θ***_***l***_ ∼ *Dir*(***β***)
>
> For each Gene (*g*)
>
> Select Gene Class: ***c***_***g***_ ∼ *Mult*(***γ***)
>
> Select Reference Group: 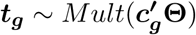
>
> For Each Sample(*j*):
>
> Select Iso Prob for Gene g: 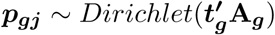
>
> Select Iso Expr: ***X***_***gj***_ ∼ *Mult*(*N*_*gj*_, ***p***_***gj***_)
>
> *N*_*gj*_ - gene expression for gene *g* of samples *j* (assumed known)

## 5 Variational Bayes Estimation and Implementation

To estimate the latent parameters (the reference group *t*, the gene class *c*, the reference group probabilities per class Θ, and the gene class probabilities ***γ***), we implement a variational Bayesian approach in the same manner as LDA, but accounting for our added structure of gene classes. First, we calculate the full joint likelihood of our model by multiplying the conditional probability distributions as shown in Equation 1. We can integrate out ***p*** to get the Dirichlet Multinomial distribution which models the isoform expression for each biological sample. This replaces *p*(***X***_***gj***_|***p***_***gj***_)*p*(***p***_***gj***_|***t***_***g***_) with *p*(***X***_***gj***_|***t***_***g***_) in our joint likelihood (Equation 1).

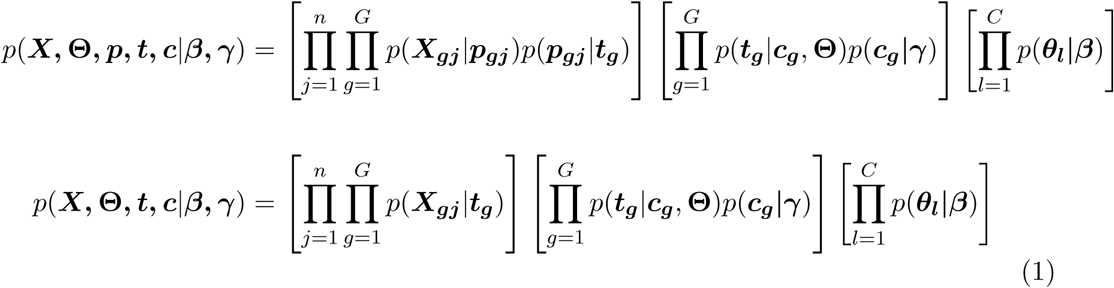

Our hope is to find *p*(**Θ, *t, c***|***X, β, γ***); however this requires the calculation of *p*(***X***|***β***), which is intractable. We instead look to find a “candidate” distribution for our full likelihood and try to estimate the parameters of this distribution which allows the candidate and full likelihoods to be similar. This procedure is known as variational inference (Jordan *and others* (1999)) and the full derivation for this can be seen in Supplementary Methods 2.1.

The candidate distribution must be carefully chosen so that it accurately depicts the full joint likelihood. This is typically handled by removing the dependence across latent variables, but maintaining the distribution proposed for each variable. We set our candidate distributions for the reference group probability vector specific to gene class (*θ*_*l*_) to be Dirichlet with a different parameter for each gene class (*η*_*l*_). The reference group membership for a given gene (*t*_*g*_) is modeled as a Multinomial distribution with a separate probability vector for each gene (*ϕ*_*g*_). The gene class membership for a given gene (*c*_*g*_) is modeled by a Multinomial distribution with a separate probability vector for each gene (*λ*_*g*_). Finally, the probability vector for gene class membership (***γ***) is modeled by the Dirichlet distribution (*ρ*). These random variables are all assumed to be independent. A diagram of these distributions is shown in figure Figure 2 b and the variational likelihood equation is given in Equation 2.

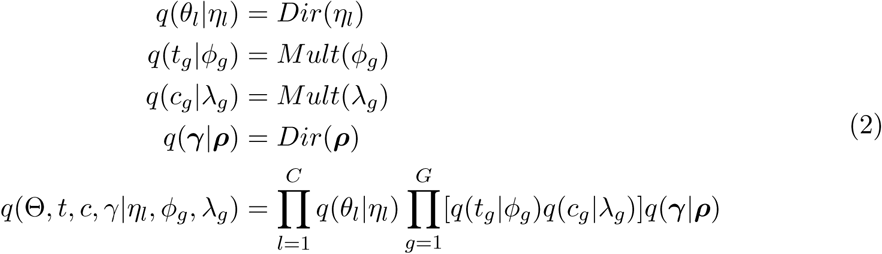

With a full joint likelihood and a candidate likelihood, the variational lower bound can be constructed by taking the difference in the expectation of both distributions (*E*_*q*_[log(*p*(***X*, Θ, *t, c***|***β, γ***))] - *E*_*q*_[log(*q*(**Θ, *t, c***))]). The expectation is taken with respect to the candidate distributions, which will remove all unobserved random variables from our estimating equation. The final result gives an equation dependent on *ϕ, η, λ, ρ, β*, and *ω* and is often called the evidence lower bound (ELBO). Equation 3 provides a formula for the ELBO and the full expression as well as its derivation can be seen in Supplementary Methods 2.1.

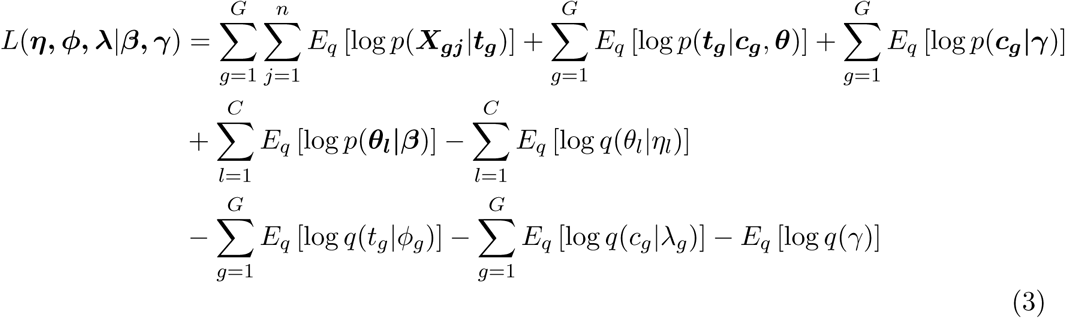

We now maximize the variational lower bound given in equation 3. Standard procedures for maximizing a multivariable distribution apply, and derivatives are calculated with respect to each variable in question. See Supplementary Methods 2.2 - 2.6 for a complete derivation of the estimates that maximize the variational lower bound. The resulting derivations give closed form solutions for *ϕ, λ, η*, and *ρ* and a Newton-Raphson algorithm for the estimation of *β* and *ω*. Closed form estimates are given in equation 4.

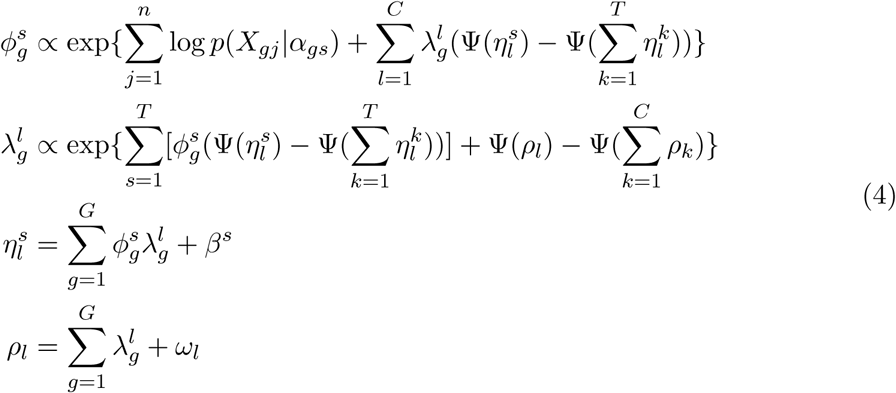

ACTOR and its estimation procedure as described above is implemented in the R package *actor*. To summarize its usage, *actor* takes as input the isoform-level estimated counts for all samples in an experimental dataset and utilizes the pre-computed Dirichlet estimates for a large, tissue-specific RNA-seq expression dataset (the GTEx project dataset, described below) as a reference panel stored internally in the R package. *actor* provides posterior estimates for all latent variables in the model as well as any additional parameters needed for fitting the model. Graphical visualizations of the posterior estimates are also available. In particular, posterior estimates of *t*_*g*_, the *T* dimensional vector for gene level membership to the reference group (*ϕ*_*g*_), posterior estimates of *c*_*g*_, the *C* dimensional vector for gene level membership to the gene class (*λ*_*g*_), and posterior estimates of *θ*_*l*_, the *T* dimensional vector of gene class estimates (*η*_*l*_) are of most interest. Detailed package vignettes provide guidance on the calculation and interpretation of these results.

## 6 Results

### 6.1 GTEx and Data Processing

We utilized data from version 7 of the Genotype-Tissue Expression project (GTEx) as a reference panel (Common Fund of the Office of the Director of the National Institutes of Health and NCI and NHGRI and NHLBI and NIDA and NIMH and NINDS). The GTEx consortium was established to study tissue-specific gene expression and identify expression quantitative trait loci (eQTL) across tissues. The dataset consists of gene expression data from 53 tissue types for over 15,000 samples collected from non-diseased individuals post-mortem. The GTEx project was supported by the Common Fund of the Office of the Director of the National Institutes of Health, and by NCI, NHGRI, NHLBI, NIDA, NIMH, and NINDS. The data used for the analyses described in this manuscript were obtained from dbGaP accession number phs000424.v7.p2 on June 30, 2017. GTEx provides a natural fit for our proposed model as the gene expression data is publicly available and the discrete groupings of tissues define the reference groups of comparison. Small sample sizes could create unreliable parameter estimates and thus tissues with less than 50 samples were removed from the analysis. Samples from the bladder (n=11), endocervix (n=6), ectocervix (n=5), fallopian tube (n=7), and kidney (n=45) were excluded. Gene filtering and parameter estimates were performed separately for each tissue. Genes that did not have at least 70% of samples with a gene count greater than 10 were not estimated and were considered not expressed for that tissue. Additionally, we considered the number of isoforms per gene as a potential filter. Increasing uncertainty in estimation of isoform expression levels and higher error rate in differential testing has been observed with increasing number of isoforms for a gene (Leng *and others* (2013); Soneson *and others* (2016)), and so we restricted our analyses to genes with five or fewer annotated isoforms. For the genes that passed filtering (8718 genes), tissue specific maximum likelihood parameter estimates were calculated for the Dirichlet-Multinomial distribution using the R package *DirichletMultinomial* (Morgan (2019)). Maximum likelihood estimates for the precision parameter of the Dirichlet Multinomial can yield very large values that can lead to small likelihoods for samples that are only slightly off from the spike. To combat this, we set the precision parameter to 50 if the calculated precision estimate was over 50.

### 6.2 Simulated Data

We utilized GTEx as our reference panel and simulated data from five GTEx tissues to demonstrate the fit of our model. Genes from five GTEx tissues (cerebellum, liver, spinal cord, pancreas, and the tibial nerve) were chosen because these tissues have been known to have the most distinct alternative splicing patterns (Yeo *and others* (2004)). A Hellinger distance was calculated on the mean isoform proportion and genes were selected to be simulated for each tissue if the Hellinger distance to all other tissues was greater than 0.1. Gene expression was simulated using a Negative Binomial distribution with *size* = 20 and *µ* = 200. Conditional on gene expression, isoform expression was allocated per gene following the Dirichlet Multinomial distribution with parameters estimated from GTEx as outlined in Section 6.1. Isoform expression was simulated for 50 samples with 100 genes simulated from cerebellum, 100 genes simulated from liver, 50 genes simulated from spinal cord, 50 genes simulated from tibial nerve, and 25 genes simulated from pancreas. Additionally, 25 genes were simulated as a mixture of the two brain tissues. These genes were selected based on having a small Hellinger distance between the two brain tissues and a large distance to all other tissues. The Dirichlet parameters for these 25 genes were then randomly chosen as coming from either cerebellum or spinal cord.

We subset to only the five tissues of interest and fit the model using ten gene classes. Ten random initializations were run to remove the possibility of the model being stuck at a local maximum. The model with the highest likelihood was chosen as the final model. ACTOR removes genes that align poorly with all tissues. Of the 375 genes that were given to the model, 369 were remaining after the model was fit. Figure 3 a provides posterior estimates for the probability vector for tissue membership (*ϕ*). Columns correspond to genes with the column annotation bars corresponding to the simulated tissue and the gene class identified by the model. Additionally, Table 1 contains Dirichlet estimates (*η*) for the gene class specific tissue probability vector. Gene classes 5-10 were empty and therefore were not shown in the table. In both the figure and table, we accurately recapture almost all of the tissue assignments that were simulated. Gene classes were also correctly identified and gene class 4 contains the genes that are uniquely cerebellum, uniquely spinal cord, and a mixture of the two. Of the 100 genes that were simulated as coming from liver, only one gene was incorrectly assigned to the brain gene class. Of the 44 genes that passed the model filtering and were assigned to be a mixture of the two brain tissues, only two of them were incorrectly classified with one being assigned to the pancreas gene class and the other to the tibial nerve gene class. Overall only 0.8% of genes were incorrectly classified.

**Table 1:** Posterior Dirichlet estimates (*η*) for the gene class specific tissue probability vector. Gene Class 4 combined both brain tissues into one group.

| Tissue | Gene Class 1 | Gene Class 2 | Gene Class 3 | Gene Class 4 |
| --- | --- | --- | --- | --- |
| Cerebellum | 0.04 | 0.04 | 0.04 | 123.94 |
| Liver | 0.04 | 99.04 | 0.04 | 0.04 |
| Pancreas | 26.04 | 0.04 | 0.04 | 0.04 |
| Tibial Nerve | 0.04 | 0.04 | 51.04 | 0.04 |
| Spinal Cord | 0.04 | 0.04 | 0.04 | 69.14 |

**Figure 3:**
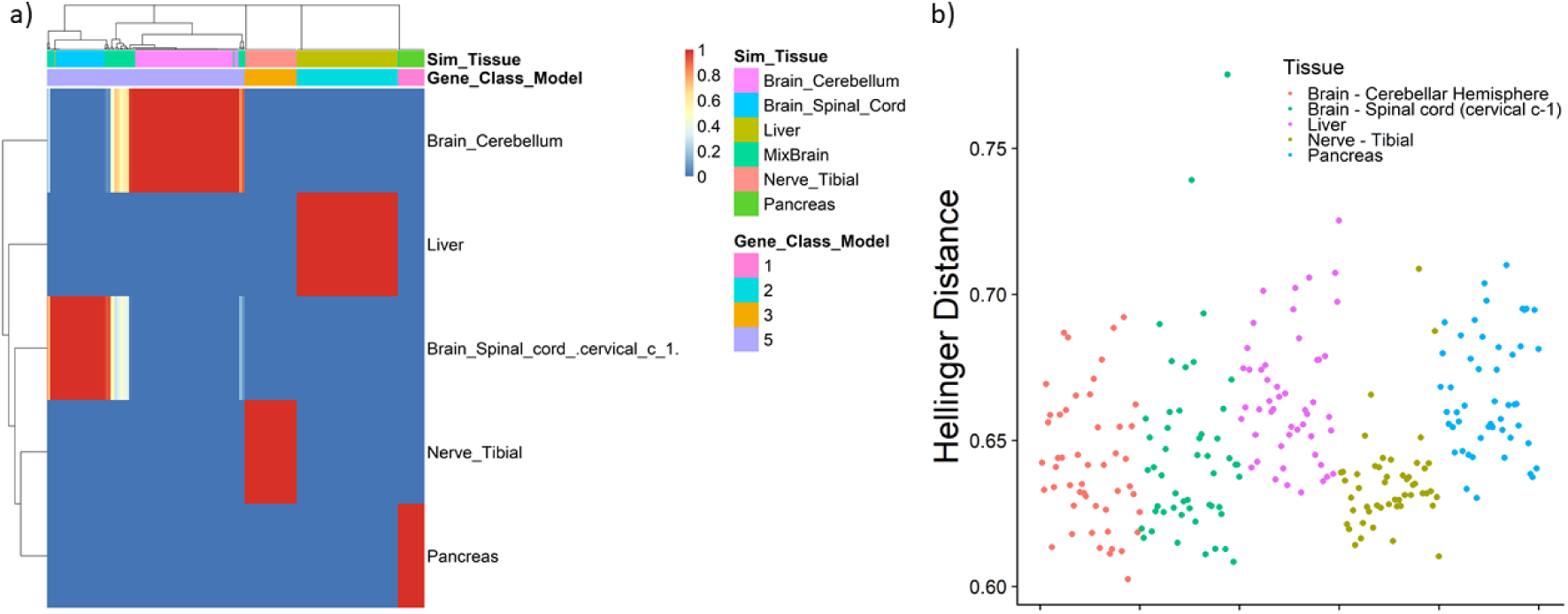
a) Multinomial posterior estimates for the probability vector for tissue membership (*ϕ*) using a simulated dataset of 50 samples fit with ten gene classes. Annotation bars above the figure represent the tissue that was simulated and the gene class that was identified by the model. Columns are genes. b) Hellinger distance between a random sample of GTEx and a simulated experimental dataset. Points are for each GTEx sample and correspond to the average Hellinger distance between that sample and each experimental sample.

Hellinger distances were calculated between each simulated sample and a subset of GTEx samples consisting of 100 samples per tissue group. Only GTEx samples from the cerebellum, liver, pancreas, tibial nerve, and spinal cord were used to allow for comparisons with the reference based model. Distances were calculated using the same set of genes that were used in the reference based model above. The average distance from simulated samples to GTEx samples was calculated, and the average distance taken over the simulated samples (Figure 3 b). It is difficult to determine which tissues were most closely aligned with the experimental samples. The results of this analysis did not include a gene-based interpretation and one possible interpretation of this analysis is that the experimental dataset did not align with liver or pancreas. This was likely due to the small number of genes that were aligned to these tissues. This demonstrates that a probabilistic model may better capture the per gene alignment as well as providing more concrete conclusions regarding splicing similarity.

The previous simulation involved simulating data using parametric assumptions with parameters informed by GTEx data: Negative Binomial total gene expression and Dirichlet Multinomial isoform expression. An additional simulation was conducted using GTEx isoform expression, combining data from two tissues to form a hybrid simulated sample. Genes that had between two and five expressed isoforms were included in this analysis and half of the genes were randomly assigned to be spleen and other half to liver, which informed the generation of isoform expression values for the simulated samples. Expression for these genes was obtained by randomly selecting the isoform expression directly from 100 GTEx samples from the corresponding tissue. In this setting, expression is not simulated parametrically as done in the first simulation, but rather as sample mixtures of existing GTEx samples. Variances were calculated for the proportion estimates of each isoform and a gene was removed from the analysis if the sum of isoform proportion variances was below 0.001 to remove genes that were similar to all GTEx tissues. For each tissue, the summation of posterior probabilities across all genes was calculated. The tissue with the smallest posterior probability sum was removed from the analysis. This process was continued until only five tissues remained. This was done to reduce the dimension of our model. After these filters were done, a total of 1975 genes remained. The model was fit using ten gene classes and ten random initializations. Figure 4 a provides the posterior estimates for the probability vector for tissue membership. Shown by the annotation bars above the figure, the model correctly identified all genes to be of the correct tissue. Table 2 provides the Dirichlet estimates for four of the ten gene classes. The six remaining gene class were empty and thus are not shown in this table. The model identified two non-empty gene classes that correspond to genes that aligned to spleen and liver, gene class 2 and 3 respectively. Only five of the 818 genes that were simulated to be liver were incorrectly classified while only two of the 1157 genes that were simulated to be spleen were incorrectly classified. Overall only 0.4% of genes were incorrectly classified.

**Table 2:**
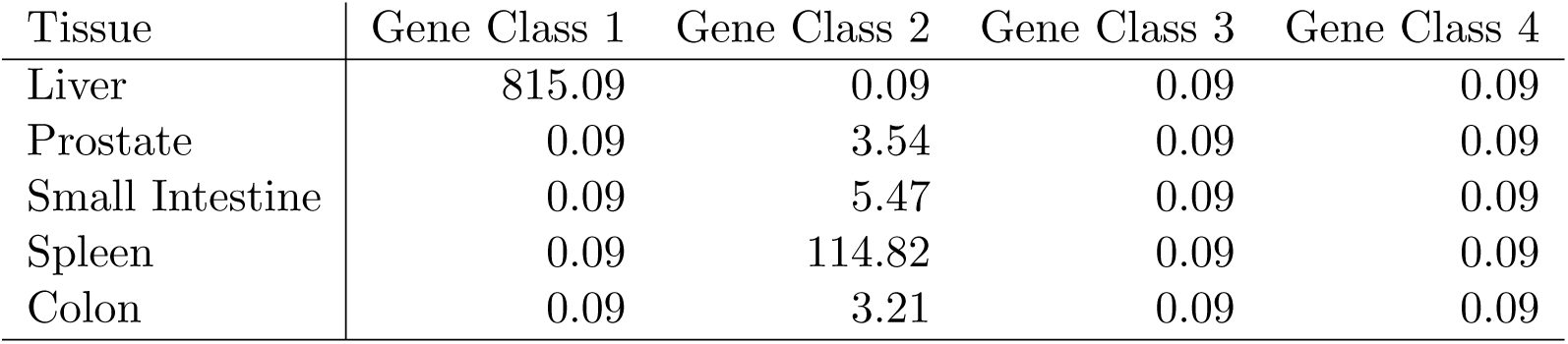
Posterior Dirichlet estimates (*η*) for the gene class specific tissue probability vector. Gene class 2 contains all genes that aligned to spleen, while gene class 3 contains all genes that aligned to liver. Gene classes 5-10 were empty and thus are not shown.

**Figure 4:**
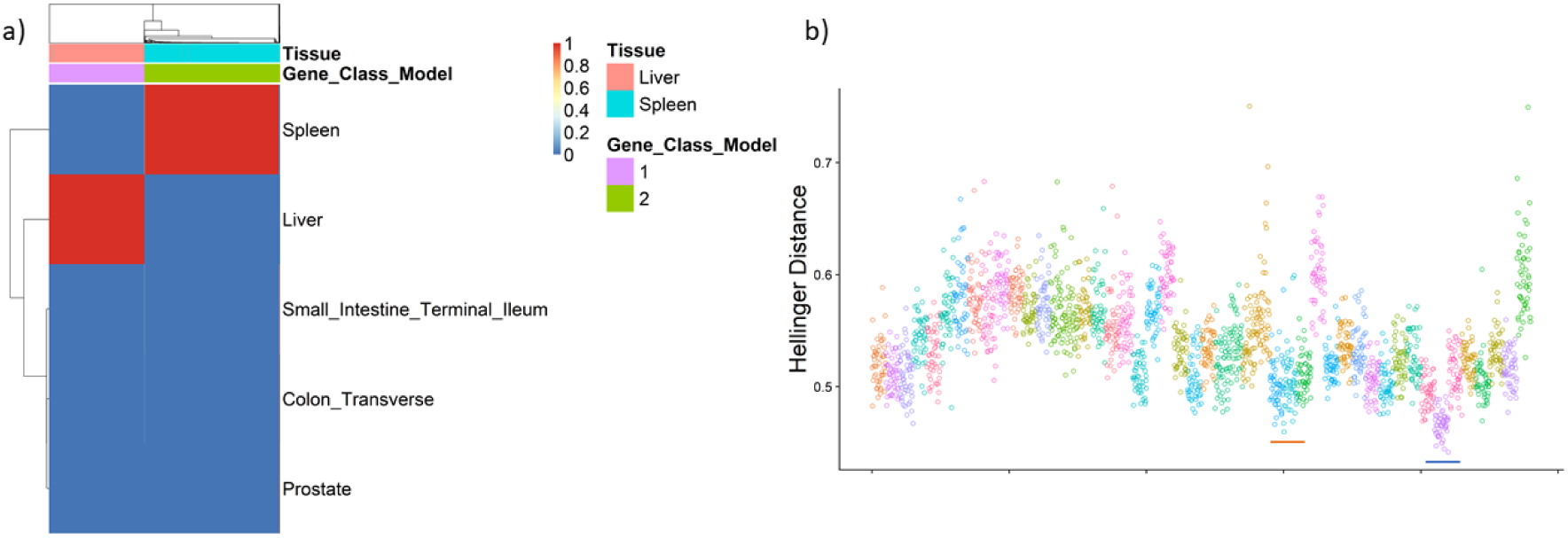
a) Posterior estimates for the probability vector for tissue membership (*ϕ*) using a dataset of 100 samples generated from a mixture of GTEx liver and spleen tissues. Annotation bars above correspond to the simulated tissue and the gene class identified by the model. b) Hellinger distance between a random sample of GTEx and a simulated experimental dataset containing spleen and liver genes. Points are for each GTEx sample and correspond to the average Hellinger distance between that sample and each experimental sample. The red bar identifies the liver GTEx samples and the blue bar identifies the spleen GTEx samples.

Hellinger distances were again calculated using the same GTEx samples. Only genes that were included in the reference panel model were included. Figure 4 b provides the average distances. A full version of this plot with an attached legend is available in Supplementary Figure 2. Spleen is correctly identified as a top tissue, however it does not appear that liver aligns well with this experimental dataset. This could be explained by the fact that liver cells exhibit some of the most distinct splicing patterns, relative to the other GTEx tissues. It is possible that genes that were not simulated to be liver had vastly different splicing patterns and thus resulted in a very large distance with all liver GTEx samples. When aggregating across genes, it is possible that liver would no longer appear similar to the experimental dataset as the larger distances would outweigh the smaller ones. It is important to note that we have constructed hybrid samples that do not naturally appear in GTEx. This will inherently provide challenges for the distance based approach and will favor our reference based model.

### 6.3 Spinal Cord RNA-seq Data

Isoform expression from an experiment analyzing lumbar spinal cord sections was analyzed using our LDA model (Batra *and others* (2016)). The isoform expression for 8 non-ALS patients was quantified using *Salmon* (Patro *and others* (2017)) with Gencode 19 to ensure that the quantified transcripts matched the set from GTEx. Only genes with two to five annotated isoforms were included for this analysis. The top 5 tissues were selected in the same manner as outlined in the previous simulation. A large number of genes were not similar to any GTEx tissue. This resulted in the strong selection of tissues with the largest variance rather than biologically accurate tissues. To combat this, genes were removed if they had an average log-likelihood across the samples less than −10. The requirement of this filtering presents a limitation of the current model and future work could address this issue. The model was fit using ten gene classes. Multiple gene classes were not found in the analysis of the spinal cord dataset. Our model aligned a large portion of the genes to the brain regions, amygdala and cerebellar, which was expected given the nerve composition of the lumbar spinal cord sections (Figure 5a). Supplementary Figures 3 and 4 provide heatmaps of individual genes that align to the amygdala and cerebellar respectively. Supplementary Figures 5-10 provide similar heatmaps for genes that aligned to liver, heart, and muscle. These heatmaps demonstrate that none of the reference panel groups are similar to the experimental dataset and that a decision is being made based on the reference group with a large variability. To ensure that the results are not driven by gene expression, a heatmap of the median gene expression for the GTEx tissues is given in Supplementary Figure 11. This heatmap is column and row ordered identically to Figure 5a and the lack of a pattern similar to the one found in Figure 5a reinforces the validity of our results.

**Figure 5:**
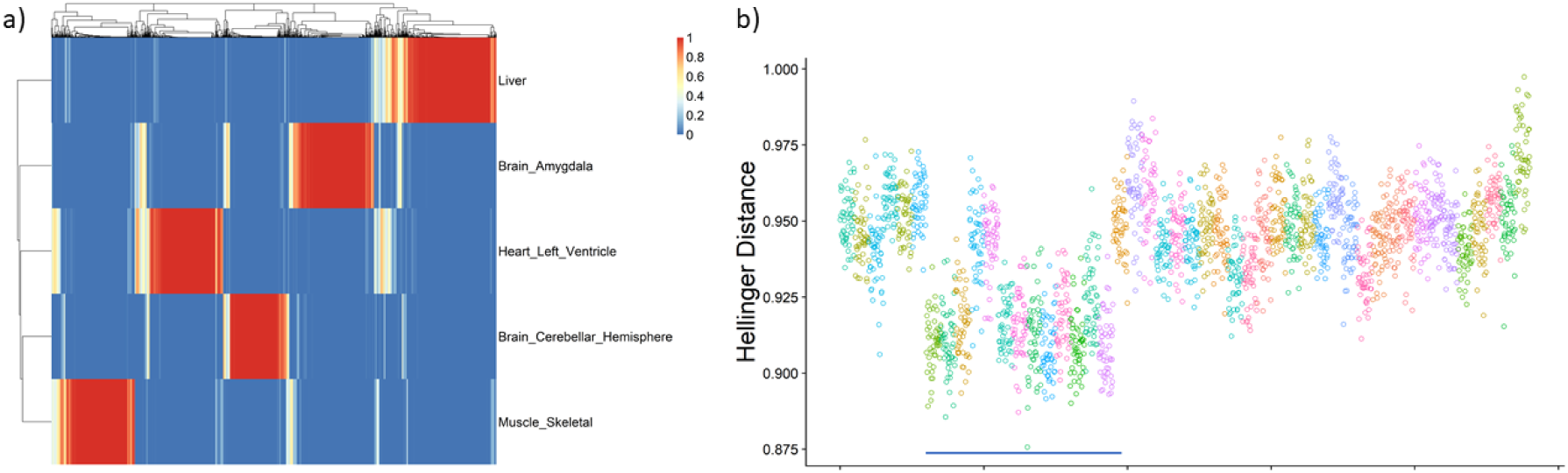
a): Posterior estimates for the probability vector for tissue membership (*ϕ*) using a real dataset of 8 motor neuron samples. b): Hellinger distance between a random sample of GTEx and motor neuron samples. Points are for each GTEx sample and correspond to the average Hellinger distance between that sample and each experimental sample. The blue bar identifies the brain GTEx samples.

Hellinger distances were calculated between each spinal cord sample and GTEx samples, with 100 samples selected per tissue as before. The average distance was calculated (Figure 5b). A full version of this plot with an attached legend is available in Supplementary Figure 12. Brain samples consistently provided small distances which would suggest that our experimental samples are most similar to brain. This is as expected, however this analysis does not provide a gene level interpretation and thus it is unknown if the alignment to brain is common across all genes or merely a subset.

## 7 Discussion

We have constructed a probabilistic model similar to LDA to compare one or more samples associated with a high dimensional vector of features — here the isoform expression patterns of RNA-seq samples — to groups in a independent reference panel. We demonstrated the effectiveness of our model through simulations constructed from the GTEx project dataset, and using Dirichlet Multinomial distributions defined by GTEx tissues as a reference panel. Our model correctly identified genes and gene classes for which isoform splicing was similar across tissues in two simulation settings. While our model was able to accurately recapture the simulated tissue, Hellinger distances were shown to only provide summary level interpretations. We showed qualitatively how distances could identify similar tissues but were not able to identify gene specific similarities.

In our evaluations, the model performs well in situations where the gene classes have orthogonal tissue probability vectors. However, our simulations involved generating hybrid samples as a proof of concept which could overly favor our method. Additionally, we only looked at genes with a small set of isoforms and implemented filtering thresholds for both genes and tissues. Future work could address these concerns to utilize a larger proportion of the data and to provide more statistically rigorous ways of removing genes. If the number of gene classes is over specified, the method will output empty gene classes however it is important to not underestimate the number of gene classes or the method will collapse all of the gene classes into one. Future work could address this issue and determine a more rigorous way of identifying the number of gene classes needed.

In our analysis of motor neuron samples comparing to GTEx, genes that were similar to all GTEx tissues or similar to none of the GTEx tissues aligned to tissues with the largest variance. This presents potential issues when utilizing an experimental dataset and reference panel that are not similar. In a real dataset, our model failed to identify gene classes. GTEx samples are collected post-mortem, while the spinal cord samples were extracted from living donors. It is possible that RNA degradation may affect quantification of isoform expression that confounded the alignment of the spinal cord samples to GTEx tissues that occurred when Dirichlet Multinomial likelihoods were computed with the fixed parameters estimated on GTEx samples. Other reference panels could be used to examine this claim. Additionally, it is possible that larger sample sizes could help in the estimation of gene classes.

Our model can also be used as a way to describe functional changes for genes that were found to be differentially spliced. After a DTU analysis, the genes with evidence for DTU can be passed to our model, fitting posterior probabilities separately for the treatment and control groups. Results from the two models could be compared to identify if the isoform switching also led to changes in the alignment of sample sets to reference groups. Our model could be extended into a full differential analysis of sample sets with respect to reference panels. Here our model was fit using a variational Bayes procedure for maximum efficiency, however a Gibbs sampling procedure may provide more stable results at the trade-off of computation time. The R package *actor* also allows for the inclusion of other reference panels and takes as input a matrix of precomputed Dirichlet estimates or a matrix of precomputed log-likelihoods.

## 8 Software Availability

ACTOR is implemented as an R package *actor* which is publicly available at https://github.com/mccabes292/actor. All of the scripts used in the analyses presented in this manuscript can be downloaded from the following respository: https://github.com/mccabes292/actorPaper.

