## Supplementary Materials for "ACTOR: a latent Dirichlet model to compare expressed isoform proportions to a reference panel"

### ACTOR - Supplementary Materials

Sean McCabe

November 22, 2019

#### 1 Supplementary Tables

| | |
|---|---|
| $j \in \{1, ..., n\}$ | sample index |
| $g \in \{1, ..., G\}$ | gene index |
| $l \in \{1, ..., C\}$ | gene class index |
| $s \in \{1, ..., T\}$ | reference group index |
| $i \in \{1, ..., I_g\}$ | isoform index for gene $g$ |
| $\omega$ | Dirichlet parameter for gene class probability vector of length $C$ |
| $\gamma$ | gene class probability vector of length $C$ |
| $c_g$ | gene class membership for gene $g$ (vector of length $T$) with a 1 in the position for the designated gene class and 0 else) |
| $\beta$ | Dirichlet parameter for the reference group probability vector of length $T$ |
| $\theta_l$ | reference group probability vector of length $T$ for gene class $l$ |
| $t_g$ | reference group membership for gene $g$ of length $T$ |
| $\alpha_{gs}$ | precomputed Dirichlet estimate for gene $g$ of the $s$ reference group of length $I_g$ |
| $p_{gj}$ | isoform probability vector for sample $j$ of gene $g$ of length $I_g$ |
| $N_{gj}$ | total gene expression for sample $j$ of gene $g$ |
| $X_{gj}$ | isoform expression for sample $j$ of gene $g$ of length $I_g$ |
| $\eta_l$ | Dirichlet parameter for the variational distribution of the reference group probability of length $T$ vector for gene class $l$ ($\theta_l$) |
| $\phi_g$ | reference group probability vector of length $T$ for gene $g$ of the variational distribution |
| $\lambda_g$ | gene class probability vector of length $C$ for gene $g$ of the variational distribution |
| $\rho$ | Dirichlet parameter of length $C$ for the gene class probability vector ($\gamma$) |

Supplementary Table 1: Notation

### 2 Supplementary Methods

#### 2.1 Variational Inference

Here we describe the derivation of the variational inference estimation procedure. From Section 5, our hope is to find $p(\boldsymbol{\theta}, \boldsymbol{t}, \boldsymbol{c}|\boldsymbol{X}, \boldsymbol{\beta}, \boldsymbol{\gamma})$; however this requires the calculation of $p(\boldsymbol{X}|\boldsymbol{\beta})$, which is intractable. We instead look to find a "candidate" distribution for our full likelihood and try to estimate the parameters of this distribution which allows the candidate and full likelihoods to be similar. This procedure is known as variational inference (**?**) and the derivation for this can be seen in Equation 1. We begin by writing out the observed log-likelihood. Here, the isoform counts ($\boldsymbol{X}$) are the only observed component of our likelihood, so we integrate or sum all other latent variables out of the full joint likelihood. We then multiply the numerator and denominator by our candidate distribution, $q(\boldsymbol{\theta}, \boldsymbol{t}, \boldsymbol{c})$, and write the likelihood as an expectation with respect to our candidate distribution $q$. Finally, we apply Jensen's Inequality to move the *log* inside of the expectation and reduce to the difference in the expectation of our full log-likelihood with respect to our candidate distribution and the expectation of our candidate distribution. This is often called the variational lower bound or evidence lower bound (ELBO) and maximizing this lower bound is equivalent to minimizing the Kullback-Leibler divergence between the full and candidate likelihoods.

$$
\begin{aligned}
\log p(\boldsymbol{X}|\boldsymbol{\beta}, \boldsymbol{\gamma}) &= \log \int_\theta \sum_c \sum_t p(\boldsymbol{X}, \boldsymbol{\theta}, \boldsymbol{t}, \boldsymbol{c}|\boldsymbol{\beta}, \boldsymbol{\gamma}) d\theta \\
&= \log \int_\theta \sum_c \sum_t \frac{p(\boldsymbol{X}, \boldsymbol{\theta}, \boldsymbol{t}, \boldsymbol{c}|\boldsymbol{\beta}, \boldsymbol{\gamma}) q(\boldsymbol{\theta}, \boldsymbol{t}, \boldsymbol{c})}{q(\boldsymbol{\theta}, \boldsymbol{t}, \boldsymbol{c})} d\theta \\
&\geq \int_\theta \sum_c \sum_t \log \left( \frac{p(\boldsymbol{X}, \boldsymbol{\theta}, \boldsymbol{t}, \boldsymbol{c}|\boldsymbol{\beta}, \boldsymbol{\gamma}) q(\boldsymbol{\theta}, \boldsymbol{t}, \boldsymbol{c})}{q(\boldsymbol{\theta}, \boldsymbol{t}, \boldsymbol{c})} \right) d\theta \\
&= E_q[\log(p(\boldsymbol{X}, \boldsymbol{\theta}, \boldsymbol{t}, \boldsymbol{c}|\boldsymbol{\beta}, \boldsymbol{\gamma}))] - E_q[\log(q(\boldsymbol{\theta}, \boldsymbol{t}, \boldsymbol{c}))] \\
&= L(\boldsymbol{\eta}, \boldsymbol{\phi}, \boldsymbol{\lambda}|\boldsymbol{\beta}, \boldsymbol{\gamma})
\end{aligned} \tag{1}
$$

Next we write out the full likelihood by taking the expectation as outlined above.

$$
\begin{aligned}
E_q[\log(p(\boldsymbol{X},\boldsymbol{\theta},\boldsymbol{t},\boldsymbol{c},\boldsymbol{\gamma}|\boldsymbol{\beta},\boldsymbol{\omega}))] &= \sum_{g=1}^{G}\sum_{j=1}^{n} E_q\left[\log p(\boldsymbol{X_{gj}}|\boldsymbol{t_g})\right] + \sum_{g=1}^{G} E_q\left[\log p(\boldsymbol{t_g}|\boldsymbol{c_g},\boldsymbol{\theta})\right] \\
&+ \sum_{g=1}^{G} E_q\left[\log p(\boldsymbol{c_g}|\boldsymbol{\gamma})\right] + \sum_{l=1}^{C} E_q\left[\log p(\boldsymbol{\theta_l}|\boldsymbol{\beta})\right] + E_q\left[\log p(\boldsymbol{\gamma}|\boldsymbol{\omega})\right] \\
E_q[\log(q(\boldsymbol{\theta},\boldsymbol{t},\boldsymbol{c},\boldsymbol{\gamma}))] &= \sum_{l=1}^{C} E_q\left[\log q(\theta_l|\eta_l)\right] + \sum_{g=1}^{G} E_q\left[\log q(t_g|\phi_g)\right] \\
&+ \sum_{g=1}^{G} E_q\left[\log q(c_g|\lambda_g)\right] + E_q\left[\log q(\boldsymbol{\gamma}|\boldsymbol{\rho})\right] \\
L(\boldsymbol{\eta},\boldsymbol{\phi},\boldsymbol{\lambda},\boldsymbol{\rho}|\boldsymbol{\beta},\boldsymbol{\omega}) &= \sum_{g=1}^{G}\sum_{j=1}^{n} E_q\left[\log p(\boldsymbol{X_{gj}}|\boldsymbol{t_g})\right] + \sum_{g=1}^{G} E_q\left[\log p(\boldsymbol{t_g}|\boldsymbol{c_g},\boldsymbol{\theta})\right] + \sum_{g=1}^{G} E_q\left[\log p(\boldsymbol{c_g}|\boldsymbol{\gamma})\right] \\
&+ \sum_{l=1}^{C} E_q\left[\log p(\boldsymbol{\theta_l}|\boldsymbol{\beta})\right] + E_q\left[\log p(\boldsymbol{\gamma}|\boldsymbol{\omega})\right] - \sum_{l=1}^{C} E_q\left[\log q(\theta_l|\eta_l)\right] \\
&- \sum_{g=1}^{G} E_q\left[\log q(t_g|\phi_g)\right] - \sum_{g=1}^{G} E_q\left[\log q(c_g|\lambda_g)\right] - E_q\left[\log q(\boldsymbol{\gamma}|\boldsymbol{\rho})\right]
\end{aligned} \tag{2}
$$

$$
\begin{aligned}
L(\boldsymbol{\eta},\boldsymbol{\phi},\boldsymbol{\lambda},\boldsymbol{\rho}|\boldsymbol{\beta},\boldsymbol{\omega}) = &\sum_{g=1}^{G}\sum_{j=1}^{n}\Bigg[\sum_{s=1}^{T}\phi_g^s\Big[\log(N_{gj}!\,) + \log(\Gamma(\sum_{i=1}^{I_g}(\alpha_{gs}^i))) - \log(\Gamma(N_{gj} + \sum_{i=1}^{I_g}\alpha_{gs}^i))\\
&+ \sum_{i=1}^{I_g}\log(\Gamma(X_{gj}^i + \alpha_{gs}^i)) - \sum_{i=1}^{I_g}\log(X_{gj}^i!\,) - \sum_{i=1}^{I_g}\log(\Gamma(\alpha_{gs}^i))\Big]\Bigg]\\
&+ \sum_{g=1}^{G}\sum_{l=1}^{C}\Big[\sum_{s=1}^{T}\phi_g^s\lambda_g^l(\Psi(\eta_l^s) - \Psi(\sum_{k=1}^{T}\eta_l^k)) + \lambda_g^l(\Psi(\rho_l) - \Psi(\sum_{k=1}^{C}\rho_k))\Big]\\
&+ \sum_{l=1}^{C}\Big[\log(\Gamma(\sum_{s=1}^{T}\beta^s) - \sum_{s=1}^{T}\log(\Gamma(\beta^s)) + \sum_{s=1}^{T}(\beta^s - 1)(\Psi(\eta_l^s) - \Psi(\sum_{k=1}^{T}\eta_l^k))\Big]\\
&+ \log\Gamma(\sum_{l=1}^{C}\omega_l) - \sum_{l=1}^{C}\log\Gamma(\omega_l) + \sum_{l=1}^{C}(\omega_l - 1)[\Psi(\rho_l) - \Psi(\sum_{k=1}^{C}\rho_l)]\\
&- \sum_{l=1}^{C}\Big[\log(\Gamma(\sum_{s=1}^{T}\eta_l^s)) - \sum_{s=1}^{T}\log(\Gamma(\eta_l^s)) + \sum_{s=1}^{T}(\eta_l^s - 1)(\Psi(\eta_l^s) - \Psi(\sum_{k=1}^{T}\eta_l^k))\Big]\\
&- \sum_{g=1}^{G}\Big[\sum_{s=1}^{T}\phi_g^s\log(\phi_g^s) + \sum_{l=1}^{C}\lambda_g^l\log(\lambda_g^l)\Big]\\
&- \log\Gamma(\sum_{l=1}^{C}\rho_l) - \sum_{l=1}^{C}\log\Gamma(\rho_l) + \sum_{l=1}^{C}(\rho_l - 1)[\Psi(\rho_l) - \Psi(\sum_{k=1}^{C}\rho_l)]
\end{aligned}
\tag{3}
$$

#### 2.2 Estimation of $\phi$

$$
\begin{aligned}
L_{\phi_g^s} = \phi_g^s \Bigg[ \sum_{j=1}^{n} \Big[ \log(N_{gj}!) + \log(\Gamma(\sum_{i=1}^{I_g} \alpha_{gs}^i)) - \log(\Gamma(N_{gj} + \sum_{i=1}^{I_g} \alpha_{gs}^i)) + \sum_{i=1}^{I_g} \log(\Gamma(X_{gj}^i + \alpha_{gs}^i)) \\
- \sum_{i=1}^{I_g} \log(X_{gj}^i!) - \sum_{i=1}^{I_g} \log(\Gamma(\alpha_{gs}^i))\Big] + \sum_{l=1}^{C} \lambda_g^l (\Psi(\eta_l^s) - \Psi(\sum_{k=1}^{T} \eta_l^k))\Bigg] \\
- \phi_g^s \log(\phi_g^s) + \epsilon_{g1}(\sum_{k=1}^{T} \phi_g^k - 1)
\end{aligned}
$$

$$
\frac{\partial L_{\phi_g^s}}{\partial \phi_g^s} = \sum_{j=1}^{n} \log p(X_{gj}|\alpha_{gs}) + \sum_{l=1}^{C} \lambda_g^l (\Psi(\eta_l^s) - \Psi(\sum_{k=1}^{T} \eta_l^k)) - \log(\phi_g^s) - 1 + \epsilon_{g1}
$$

$$
\phi_g^s \propto \exp\{\sum_{j=1}^{n} \log p(X_{gj}|\alpha_{gs}) + \sum_{l=1}^{C} \lambda_g^l (\Psi(\eta_l^s) - \Psi(\sum_{k=1}^{T} \eta_l^k))\} \tag{4}
$$

#### 2.3 Estimation of $\lambda$

$$
L_{\lambda_g^l} = \lambda_g^l [\sum_{s=1}^{T} \phi_g^s (\Psi(\eta_l^s) - \Psi(\sum_{k=1}^{T} \eta_l^k)) + \Psi(\rho_l) - \Psi(\sum_{k=1}^{C} \rho_k)] - \lambda_g^l \log(\lambda_g^l) + \epsilon_{g2}(\sum_{k=1}^{C} \lambda_g^k - 1)
$$

$$
\frac{\partial L_{\lambda_g^l}}{\partial \lambda_g^l} = \sum_{s=1}^{T} [\phi_g^s (\Psi(\eta_l^s) - \Psi(\sum_{k=1}^{T} \eta_l^k))] + \Psi(\rho_l) - \Psi(\sum_{k=1}^{C} \rho_k) - 1 - \log(\lambda_g^l) + \epsilon_{g2}
$$

$$
\lambda_g^l \propto \exp\{\sum_{s=1}^{T} [\phi_g^s (\Psi(\eta_l^s) - \Psi(\sum_{k=1}^{T} \eta_l^k))] + \Psi(\rho_l) - \Psi(\sum_{k=1}^{C} \rho_k)\} \tag{5}
$$

#### 2.4 Estimation of $\eta$

$$
\begin{aligned}
L_\eta &= \sum_{l=1}^{C}\sum_{s=1}^{T}\Big[\sum_{g=1}^{G}\phi_g^s\lambda_g^l + \beta^s - 1\Big](\Psi(\eta_l^s) - \Psi(\sum_{k=1}^{T}\eta_l^k)) \\
&- \sum_{l=1}^{C}\Big[\log(\Gamma(\sum_{s=1}^{T}\eta_l^s)) - \sum_{s=1}^{T}\log(\Gamma(\eta_l^s)) + \sum_{s=1}^{T}(\eta_l^s - 1)(\Psi(\eta_l^s) - \Psi(\sum_{k=1}^{T}\eta_l^k))\Big] \\
\frac{\partial L}{\partial \eta_l^s} &= \Big[\sum_{g=1}^{G}\phi_g^s\lambda_g^l + \beta^s - 1\Big]\Psi'(\eta_l^s) - \sum_{k=1}^{T}\Big[\sum_{g=1}^{G}\phi_g^k\lambda_g^l + \beta^k - 1\Big]\Psi'(\sum_{k=1}^{T}\eta_l^k) \\
&- \Psi(\sum_{k=1}^{T}\eta_l^k) + \Psi(\eta_l^s) - (\Psi(\eta_l^s) - \Psi(\sum_{k=1}^{T}\eta_l^k)) - (\eta_l^s - 1)\Psi'(\eta_l^s) + \sum_{k=1}^{T}(\eta_l^k - 1)\Psi'(\sum_{k=1}^{T}\eta_l^k) \\
&= \Psi'(\eta_l^s)\Big[\sum_{g=1}^{G}\phi_g^s\lambda_g^l + \beta^s - \eta_l^s\Big] + \sum_{k=1}^{T}\Psi'(\sum_{k=1}^{T}\eta_l^k)\left[\sum_{g=1}^{G}\phi_g^k\lambda_g^l + \beta^k - \eta_l^k\right] \\
\eta_l^s &= \sum_{g=1}^{G}\phi_g^s\lambda_g^l + \beta^s
\end{aligned}
\tag{6}
$$

#### 2.5 Estimation of $\rho$

$$L_{\rho_l} = \sum_{g=1}^{G}\sum_{l=1}^{C}\lambda_g^l[\Psi(\rho_l) - \Psi(\sum_{k=1}^{C}\rho_k)] + \sum_{l=1}^{C}(\omega_l - 1)[\Psi(\rho_l) - \Psi(\sum_{k=1}^{C}\rho_k)] - \log\Gamma(\sum_{k=1}^{C}\rho_k)$$
$$+ \sum_{l=1}^{C}\log\Gamma(\rho_l) - \sum_{l=1}^{C}(\rho_l - 1)[\Psi(\rho_l) - \Psi(\sum_{k=1}^{C}\rho_k)]$$

$$= \sum_{l=1}^{C}(\sum_{g=1}^{G}\lambda_g^l + \omega_l - 1)\Psi(\rho_l) - \sum_{l=1}^{C}(\sum_{g=1}^{G}\lambda_g^l + \omega_l - 1)\Psi(\sum_{k=1}^{C}\rho_k) - \log\Gamma(\sum_{k=1}^{C}\rho_k)$$
$$+ \sum_{l=1}^{C}\log\Gamma(\rho_l) - \sum_{l=1}^{C}(\rho_l - 1)\Psi(\rho_l) + \sum_{l=1}^{C}(\rho_l - 1)\Psi(\sum_{k=1}^{C}\rho_k)$$

$$\frac{\partial L_{\rho_l}}{\partial \rho_l} = (\sum_{g=1}^{G}\lambda_g^l + \omega_l - 1)\Psi'(\rho_l) - \sum_{k=1}^{C}(\sum_{g=1}^{G}\lambda_g^k + \omega_k - 1)\Psi'(\sum_{k=1}^{C}\rho_k) - \Psi(\sum_{k=1}^{C}\rho_k)$$
$$+ \Psi(\rho_l) - (\rho_l - 1)\Psi'(\rho_l) - \Psi(\rho_l) + \sum_{k=1}^{C}(\rho_k - 1)\Psi'(\sum_{k=1}^{C}\rho_k) + \Psi(\sum_{k=1}^{C}\rho_k)$$
$$= \Psi'(\rho_l)(\sum_{g=1}^{G}\lambda_g^l + \omega_l - \rho_l) - \Psi'(\sum_{k=1}^{C}\rho_k)\sum_{k=1}^{C}(\sum_{g=1}^{G}\lambda_g^k + \omega_k - \rho_k)$$

$$\rho_l = \sum_{g=1}^{G}\lambda_g^l + \omega_l \tag{7}$$

#### 2.6 Estimation of $\beta$

$$
\begin{aligned}
L_{\beta^s} &= \sum_{l=1}^{C}[\log(\Gamma(\sum_{k=1}^{T}\beta^k)) - \log(\Gamma(\beta^s))] + (\beta^s - 1)(\Psi(\eta_l^s) - \Psi(\sum_{k=1}^{T}\eta_l^k)) \\
\frac{\partial L_{\beta^s}}{\partial \beta^s} &= C(\Psi(\sum_{k=1}^{T}\beta_k) - \Psi(\beta^s)) + \sum_{l=1}^{C}(\Psi(\eta_l^s) - \Psi(\sum_{k=1}^{T}\eta_l^k)) \\
\frac{\partial^2 L_{\beta^2}}{\partial(\beta^s)^2} &= C(\Psi'(\sum_{k=1}^{T}\beta^k) - \Psi'(\beta^s)) \\
\frac{\partial^2 L_{\beta^2}}{\partial \beta^s \beta^u} &= C\Psi'(\sum_{k=1}^{T}\beta^k) \\
H &= diag(C(\Psi'(\beta^1)), ..., C(\Psi'(\beta^T))) + 11^T(C\Psi'(\sum_{k=1}^{T}\beta^k)) \\
&= Z + a11^T \\
H^{-1} &= Z^{-1} - \frac{Z^{-1}11^T Z^{-1}}{\frac{1}{a} + 1^T Z^{-1} 1}
\end{aligned}
\tag{8}
$$

Inverting $H$ is found by application the Sherman-Morrison inverse formula.

Estimation of $\omega$ is done similarly.

### 3 Supplementary Figures

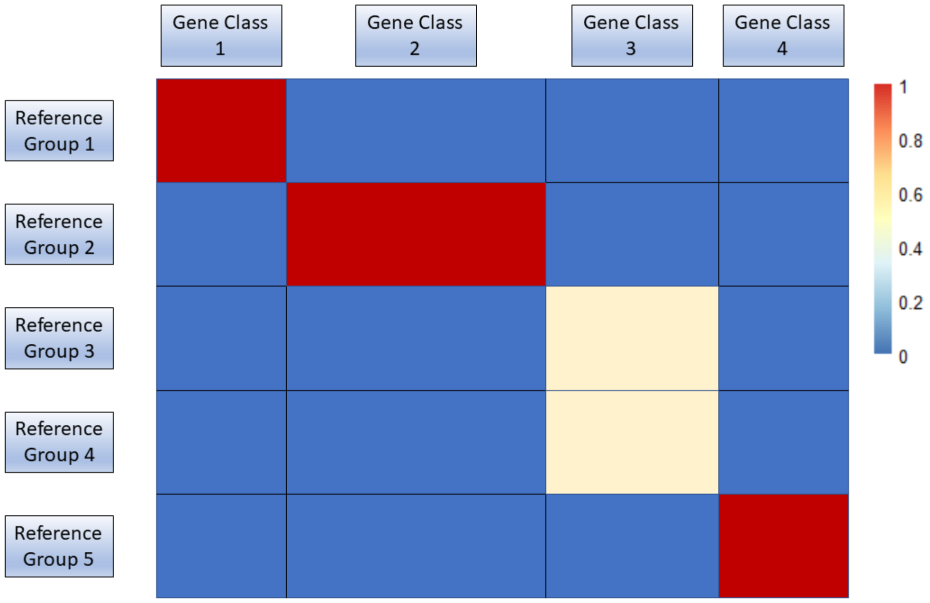


Supplementary Figure 1: Hellinger distance between a random sample of GTEx and a simulated experimental dataset containing spleen and liver genes. Points are for each GTEx sample and correspond to the average Hellinger distance between that sample and each experimental sample.

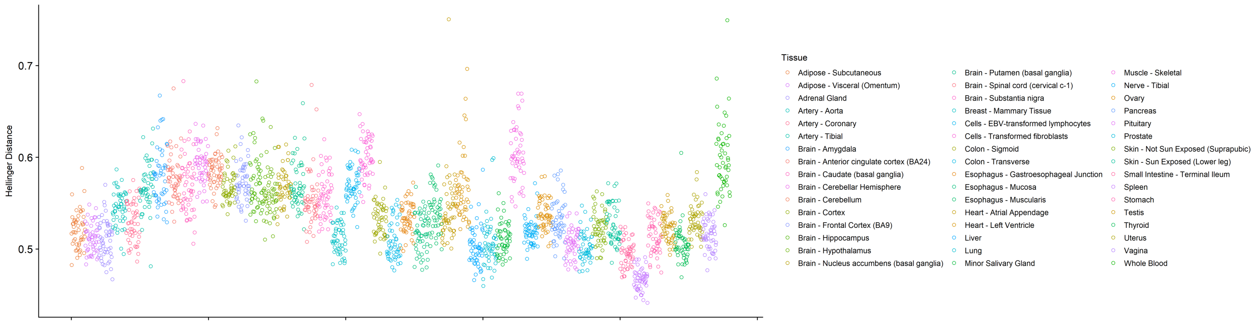


Supplementary Figure 2: Hellinger distance between a random sample of GTEx and a simulated experimental dataset containing spleen and liver genes. Points are for each GTEx sample and correspond to the average Hellinger distance between that sample and each experimental sample.

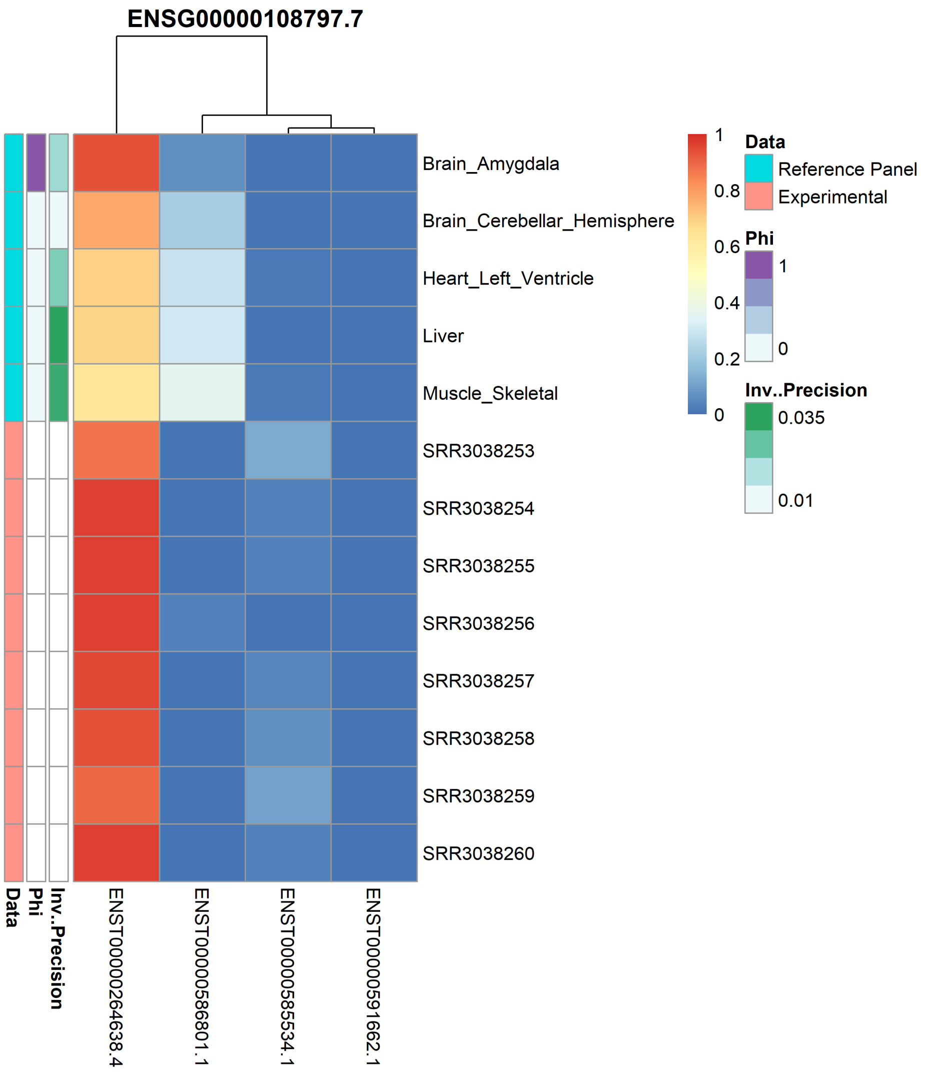


Supplementary Figure 3: Heatmap of the isoform proportions for gene ENSG00000108797.7, which aligned to amygdala. Each column corresponds to an isoform of this gene with the rows corresponding to either a tissue group from GTEx or an experimental sample indicated by the first annotation bar. Cells are the estimated isoform proportions for GTEx tissues and the observed isoform proportions for experimental samples. The second annotation bar shows the posterior estimate from the model for the tissue group membership. The last annotation bar is the inverse precision for the GTEx tissue for these gene. Experimental samples will not have a value for the last two annotation bars.

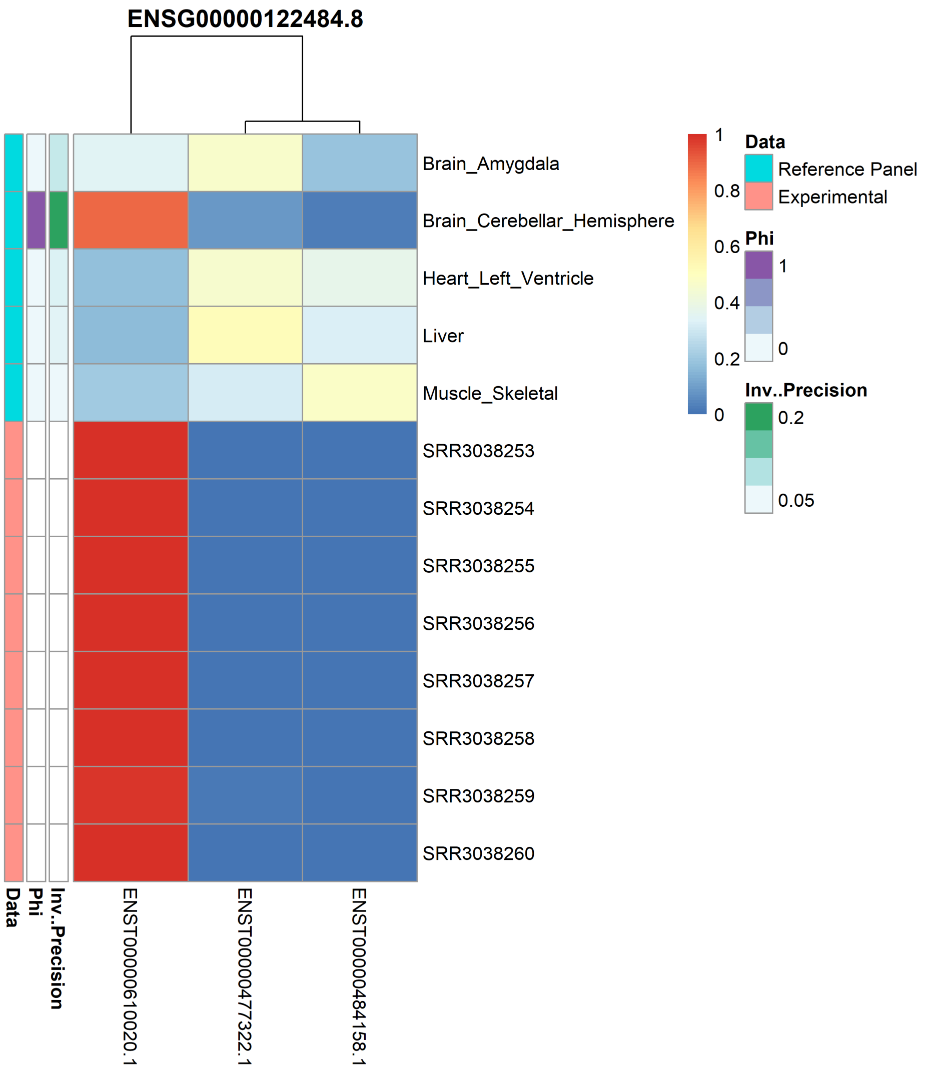


Supplementary Figure 4: Heatmap of the isoform proportions for gene ENSG00000122484.8, which aligned to cerebellar. Each column corresponds to an isoform of this gene with the rows corresponding to either a tissue group from GTEx or an experimental sample indicated by the first annotation bar. Cells are the estimated isoform proportions for GTEx tissues and the observed isoform proportions for experimental samples. The second annotation bar shows the posterior estimate from the model for the tissue group membership. The last annotation bar is the inverse precision for the GTEx tissue for these gene. Experimental samples will not have a value for the last two annotation bars.

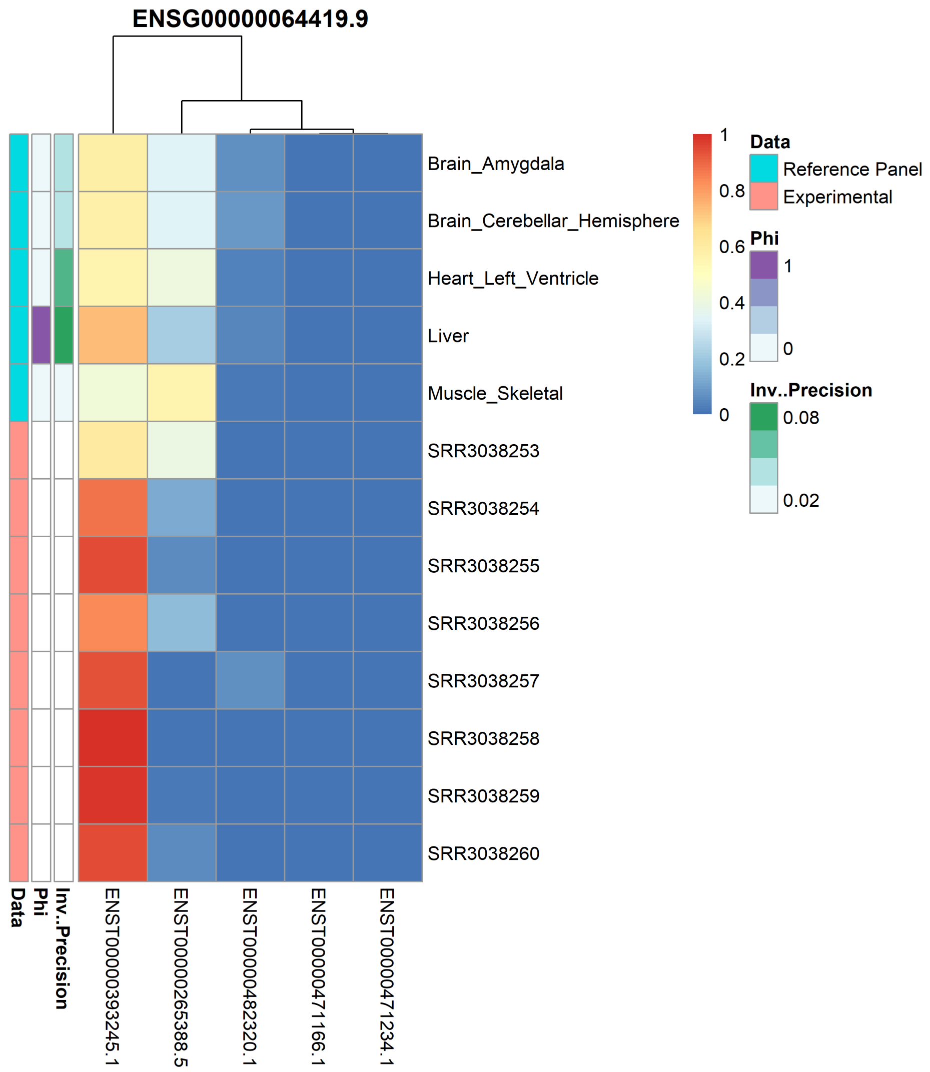


Supplementary Figure 5: Heatmap of the isoform proportions for gene ENSG00000064419.9, which aligned to liver. Each column corresponds to an isoform of this gene with the rows corresponding to either a tissue group from GTEx or an experimental sample indicated by the first annotation bar. Cells are the estimated isoform proportions for GTEx tissues and the observed isoform proportions for experimental samples. The second annotation bar shows the posterior estimate from the model for the tissue group membership. The last annotation bar is the inverse precision for the GTEx tissue for these gene. Experimental samples will not have a value for the last two annotation bars.

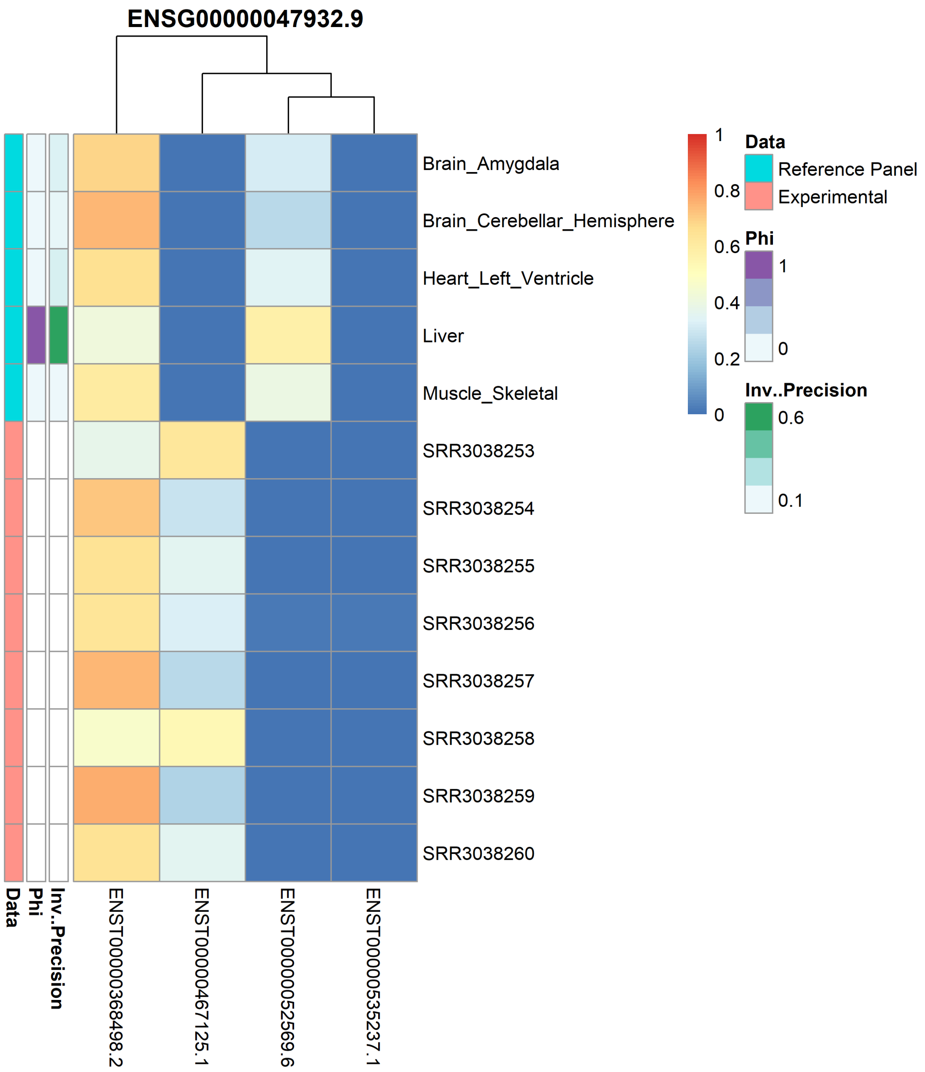


Supplementary Figure 6: Heatmap of the isoform proportions for gene ENSG00000047932.9, which aligned to liver. Each column corresponds to an isoform of this gene with the rows corresponding to either a tissue group from GTEx or an experimental sample indicated by the first annotation bar. Cells are the estimated isoform proportions for GTEx tissues and the observed isoform proportions for experimental samples. The second annotation bar shows the posterior estimate from the model for the tissue group membership. The last annotation bar is the inverse precision for the GTEx tissue for these gene. Experimental samples will not have a value for the last two annotation bars.

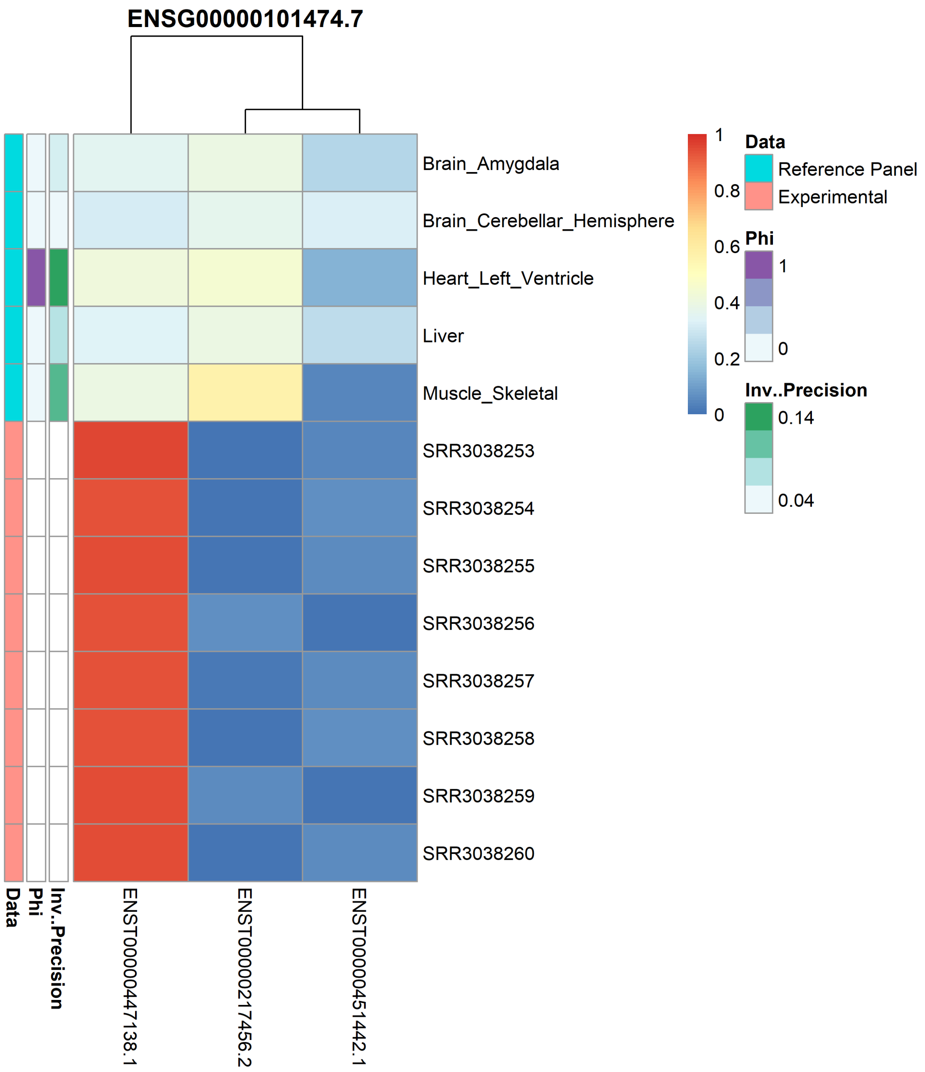


Supplementary Figure 7: Heatmap of the isoform proportions for gene ENSG00000101474.7, which aligned to heart. Each column corresponds to an isoform of this gene with the rows corresponding to either a tissue group from GTEx or an experimental sample indicated by the first annotation bar. Cells are the estimated isoform proportions for GTEx tissues and the observed isoform proportions for experimental samples. The second annotation bar shows the posterior estimate from the model for the tissue group membership. The last annotation bar is the inverse precision for the GTEx tissue for these gene. Experimental samples will not have a value for the last two annotation bars.

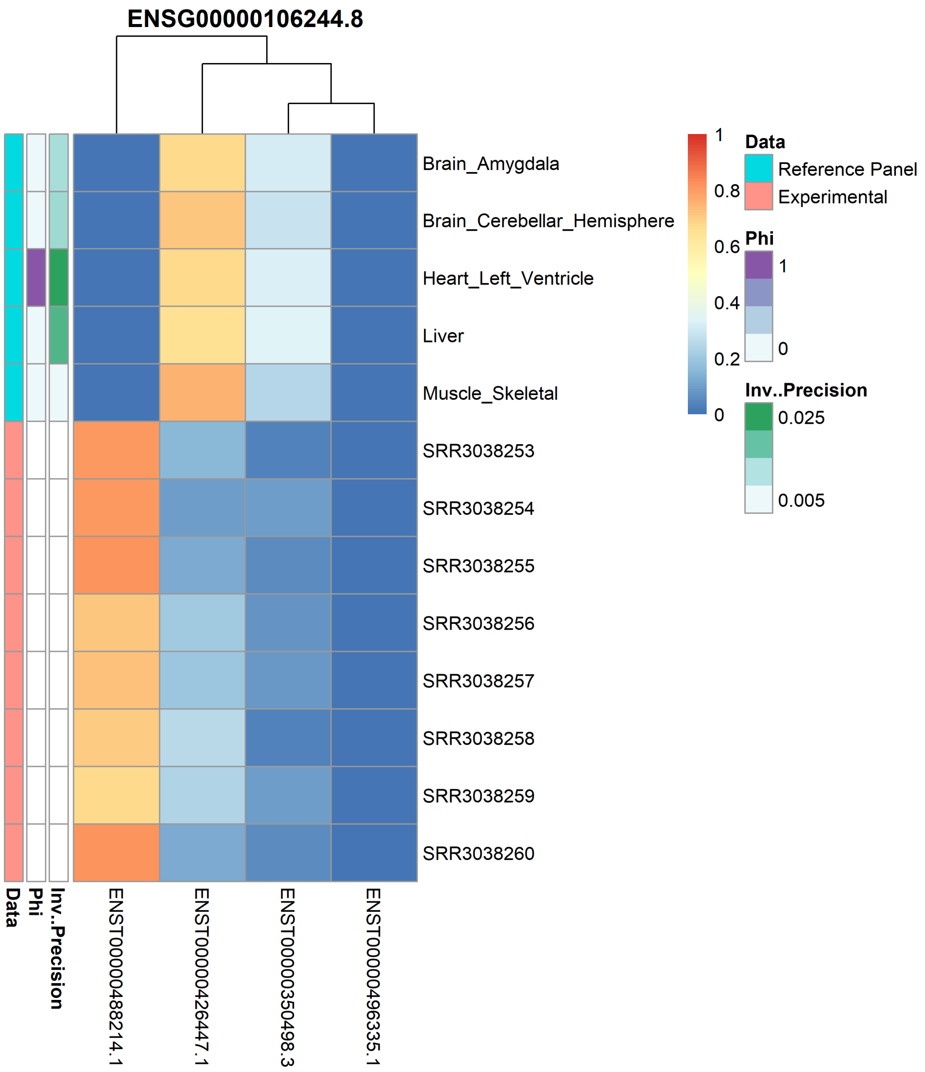


Supplementary Figure 8: Heatmap of the isoform proportions for gene ENSG00000106244.8, which aligned to heart. Each column corresponds to an isoform of this gene with the rows corresponding to either a tissue group from GTEx or an experimental sample indicated by the first annotation bar. Cells are the estimated isoform proportions for GTEx tissues and the observed isoform proportions for experimental samples. The second annotation bar shows the posterior estimate from the model for the tissue group membership. The last annotation bar is the inverse precision for the GTEx tissue for these gene. Experimental samples will not have a value for the last two annotation bars.

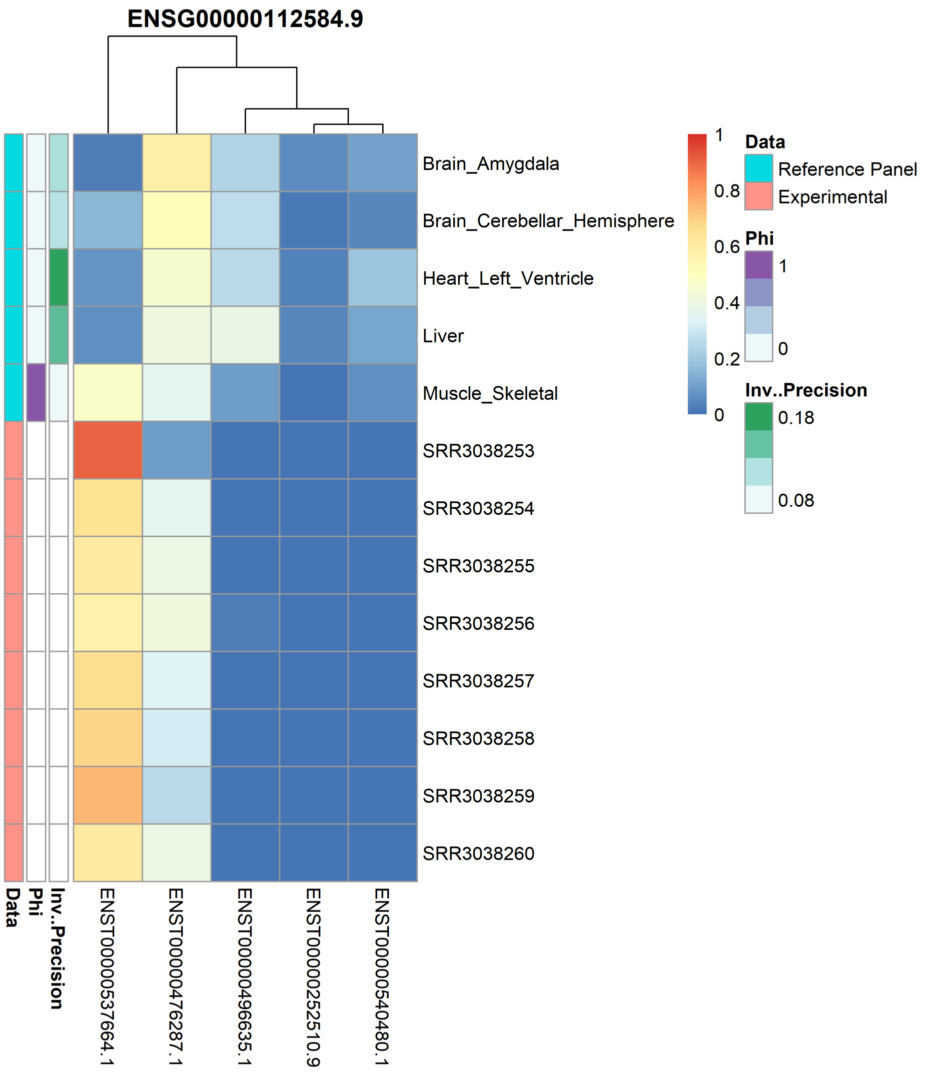


Supplementary Figure 9: Heatmap of the isoform proportions for gene ENSG00000112584.9, which aligned to muscle. Each column corresponds to an isoform of this gene with the rows corresponding to either a tissue group from GTEx or an experimental sample indicated by the first annotation bar. Cells are the estimated isoform proportions for GTEx tissues and the observed isoform proportions for experimental samples. The second annotation bar shows the posterior estimate from the model for the tissue group membership. The last annotation bar is the inverse precision for the GTEx tissue for these gene. Experimental samples will not have a value for the last two annotation bars.

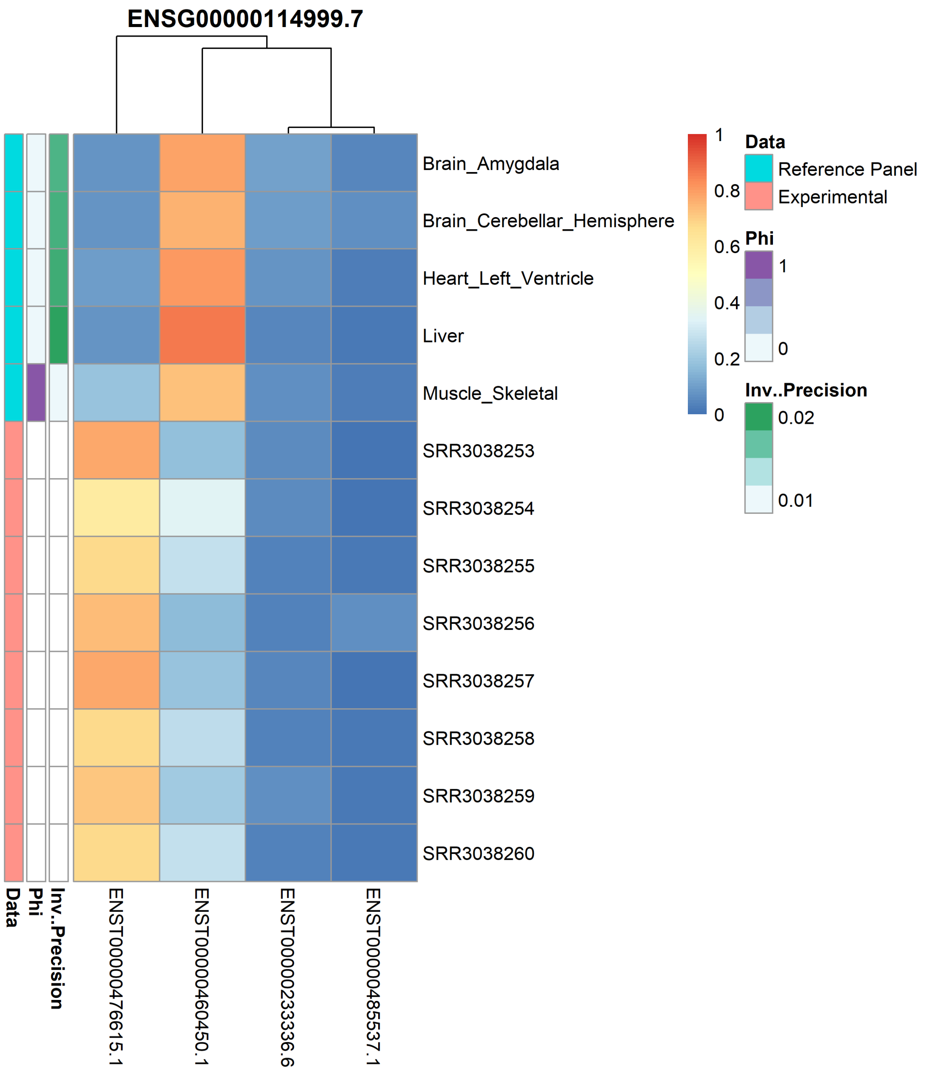


Supplementary Figure 10: Heatmap of the isoform proportions for gene ENSG00000114999.7, which aligned to muscle. Each column corresponds to an isoform of this gene with the rows corresponding to either a tissue group from GTEx or an experimental sample indicated by the first annotation bar. Cells are the estimated isoform proportions for GTEx tissues and the observed isoform proportions for experimental samples. The second annotation bar shows the posterior estimate from the model for the tissue group membership. The last annotation bar is the inverse precision for the GTEx tissue for these gene. Experimental samples will not have a value for the last two annotation bars.

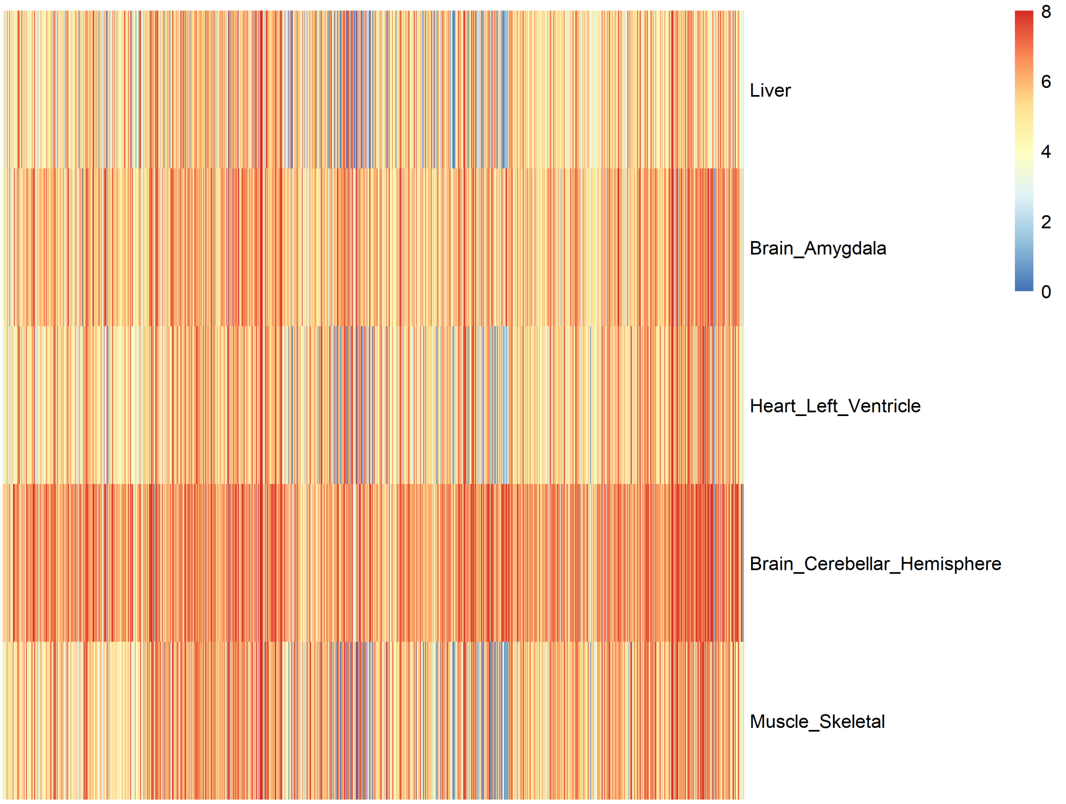


Supplementary Figure 11: Heatmap of median gene expression of each GTEx tissue used in the real data analysis. Each column is a gene and each row is a GTEx tissue. Cells are the log transformed gene expression. Rows and columns have been ordered in the same order as Figure 7 from the main text.

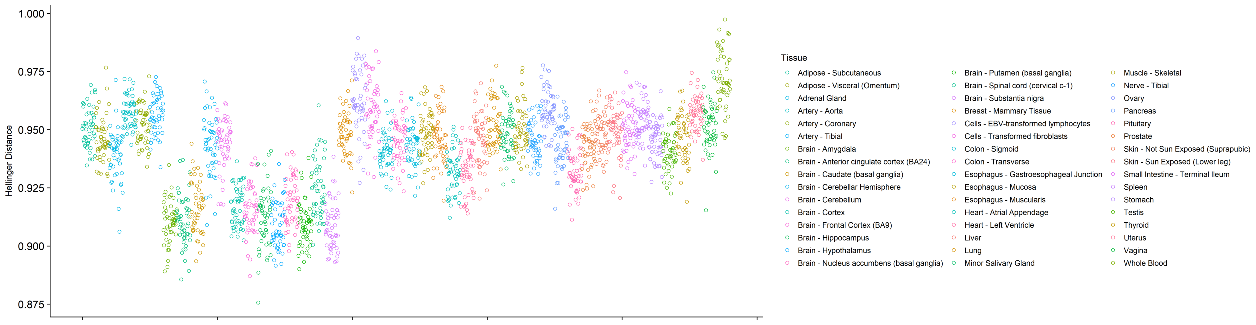


Supplementary Figure 12: Hellinger distance between a random sample of GTEx and motor neuron samples. Points are for each GTEx sample and correspond to the average Hellinger distance between that sample and each experimental sample.
